# Deciphering *APOBEC1* in Avians: Unravelling loss events and functional insights

**DOI:** 10.64898/2025.12.03.692250

**Authors:** Aswin S Soman, Nagarjun Vijay

## Abstract

In vertebrates, cytidine-to-uracil (C-to-U) editing is mediated by the *AID*/*APOBEC* family of deaminases, with *APOBEC1* (*A1*) known to catalyse precise RNA editing of apolipoprotein B (apoB) transcripts in mammals. Despite its well-characterised role in mammals, the evolutionary history and functional divergence of *A1* across birds remain underexplored. Here, we investigate the evolutionary trajectory of *A1* in birds, where both the presence of the gene and apoB RNA editing activity have been questioned. Through a comprehensive in silico analysis of 81 avian genomes, we identify recurrent disruptions and catalytic inactivation of *A1* in multiple lineages. Comparative sequence and structural analyses reveal a lack of domains and key residues essential for RNA binding, dimerisation, and cofactor interaction, suggesting a role in DNA editing. Furthermore, genome-wide screening for *A1*-associated G-to-A mutations in long terminal repeat (LTR) retrotransposons demonstrates that species with higher endogenous retrovirus (ERV) loads retain more DNA editing signatures, consistent with a defensive role of *A1* against retroelements. In contrast, species with low ERV content exhibit relaxed selection and frequent *A1* pseudogenisation. Together, these findings support the hypothesis that DNA editing represents the ancestral function of *A1*, with RNA editing in mammals evolving later as an exaptation following the expansion of *A3* and changes in retroviral pressures.

## 1) Introduction

Genetic diversity serves as the engine of evolution, driven by molecular processes that generate variation and enhance adaptability to changing environmental conditions. Among the mechanisms contributing to this variability, RNA editing stands out as a process that diversifies proteomes by altering RNA without hardwiring changes into the DNA (Gommans et al. 2009; Duan et al. 2023a, b; Qi et al. 2024). The evolutionary significance of RNA editing in adaptation has been explained through two contrasting hypotheses. The “adaptive hypothesis” posits that RNA editing serves as an adaptive mechanism to enhance phenotypic plasticity, enabling organisms to fine-tune protein functions in response to changing environments, as evidenced by the higher-than-neutral expectation prevalence of nonsynonymous editing sites in conserved genes (Garrett and Rosenthal 2012; Alon et al. 2015; Liscovitch-Brauer et al. 2017; Buchumenski et al. 2017; Duan et al. 2023a, b; Xin et al. 2023). Conversely, evidence from human studies challenges this hypothesis, showing that nonsynonymous editing sites are significantly less frequent than synonymous ones, occur at lower frequencies, and are rare in essential genes (Xu and Zhang 2014; Liu and Zhang 2018). While studying the edit sites is important, investigating the origin and evolutionary dynamics of RNA editing systems is just as crucial, as its adaptive significance is deeply linked to its evolutionary history.

RNA editing refers to the biological process by which RNA is modified co- or post-transcriptionally through the insertion, deletion or substitution of nucleotides (Farajollahi and Maas 2010). Broadly, it is part of the larger repertoire of RNA modifications, which also includes 5′ capping, 3′ polyadenylation, and splicing. In vertebrates, substitution is the primary form of RNA editing, with the most extensively studied types being the conversion of adenosine (A) to inosine (I) and cytosine (C) to uracil (U) (Maas and Rich 2000; Blanc and Davidson 2003; Wedekind et al. 2003). The conversion of A-to-I is mediated by “adenosine deaminase that acts on RNA” (*ADAR*) (Savva et al. 2012; Mannion et al. 2015) and “adenosine deaminase active on tRNA” (*ADAT*) (Rafels-Ybern et al. 2015), while C-to-U conversion is mediated by *AID*/*APOBEC* family. Unlike ADAR/ADAT proteins, some of the *AID*/*APOBEC* family members can act on both ssRNA and ssDNA. Despite these functional differences, both enzyme families share a conserved catalytic region referred to as the Zinc-Dependent Deaminase Domain (ZDD) (Harris et al. 2002; Goodman et al. 2012; Aydin et al. 2014; Shaban et al. 2016). The ZDD contains active sites critical for performing deamination, characterised by a conserved consensus amino acid motif HxEx_25-30_PCx_2-4_C, where x denotes any amino acid (Conticello et al. 2005; Iyer et al. 2011). Given the extensive gene duplications, losses, and diversification within the *AID*/*APOBEC* family, exploring its evolutionary trajectory can provide key insights into the mechanisms driving its dynamic evolution (Krishnan et al. 2018; Wang and Han 2023).

The *AID*/*APOBEC* family comprises 12 members: *AID*, *APOBEC1* (*A1*), *APOBEC2* (*A2*), seven *APOBEC3* proteins (*A3A*, *A3B*, *A3C*, *A3D*, *A3F*, *A3G*, and *A3H*), *APOBEC4* (*A4*), and *APOBEC5* (*A5*). *AID* and *A2* are the most ancestral members of the family, from which other members have evolved (Conticello et al. 2005). *A1* and *A3* are late arrivals in the family, likely arising from the duplication of *AID*. *A1* modifies the RNA of apolipoprotein B (apoB), a crucial component in lipid transportation (Hirano et al. 1997; Davidson and Shelness 2000), while A3 proteins act on RNA and ssDNA and hypermutate them to perform functions mainly related to restricting retroviruses and retrotransposons (Aydin et al. 2014; Sharma et al. 2015; Salter et al. 2016). The physiological function of *A1* is well established in placental mammals, in which the C-to-U conversion in apoB RNA leads to a change of glutamate (CAA) to a stop codon (UAA), resulting in a shorter protein called apoB48 (Driscoll and Casanova 1990; Lau et al. 1991; Daniels et al. 2009). This editing process is highly precise, targeting a single nucleotide (C6666) in the 14kb gene (Damsteegt et al. 2018). Its specificity and efficiency are attributed to two key factors: cis-acting elements (e.g., the 5’ efficiency sequence, AU-rich region, spacer element, mooring sequence, and 3’ efficiency sequence) located upstream and downstream of the target base (Shah et al. 1991; Backus and Smith 1992), and trans-acting factors such as A1CF and RBM47, which interact with A1 to form the editing complex (Mehta and Driscoll 2002; Fossat et al. 2014). Among these, the mooring sequence (6671-6681) is the primary cis-acting element, serving as the key recognition motif; mutations in this region consistently lead to a significant loss or complete disruption of RNA editing (Backus and Smith 1991; Davidson and Shelness 2000). The evolutionary origin of such exceptional substrate specificity remains unresolved. However, tracing the emergence, retention, and loss of *A1* across vertebrates and examining the molecular features that influence these events may help elucidate the evolutionary forces shaping RNA editing and its functional diversification.

*A1* was initially considered unique to mammals, given its absence in chickens, the lack of apoB RNA editing activity detected in non-mammalian vertebrates, and a surge in *A1* identification across various mammalian species (Teng and Davidson 1992; Greeve et al. 1993; Anant et al. 1998; Fujino et al. 1999; Conticello et al. 2005). Subsequent findings, however, cloned *A1* homologs in lizard and zebra finch and, in silico analyses, identified *A1* in various sauropsids and lungfish, suggesting that *A1* originated in early tetrapods (Conticello et al. 2005; Severi et al. 2011; Wang and Han 2023). Despite the widespread presence of *A1* in tetrapods, to the best of our knowledge, there is no evidence of apoB RNA editing occurring outside of mammals. Lizard *A1*, although incapable of editing apoB RNA in cell cultures, retained DNA editing, a function also observed in some mammalian *A1*, which edits the single-stranded DNA (ssDNA) of retroviruses and retrotransposons (Harris et al. 2002; Ikeda et al. 2008, 2017; Petit et al. 2009). These findings support the notion that apoB RNA editing is mammalian-specific, while DNA editing is likely *A1*’s ancestral function. In mammals, *A1* is likely co-opted for apoB RNA editing, as the emergence of *A3* in placental mammals took over antiviral functions (Münk et al. 2012; Ito et al. 2020). Compared to mammals, birds have a lower endogenous retrovirus (ERV) load, likely due to higher DNA deletion rates and strong selection against new insertions, which contribute to genome size reduction, a trait associated with their flight adaptation (Cui et al. 2014; Zheng et al. 2024). A study found that among various mechanisms, gene silencing contributes the most to ERV suppression in birds, resulting in a strong correlation between genome-wide coding gene dN/dS and ERV load, surpassing the effects of immunity and DNA recombination (Cui et al. 2014). We propose that *A1*’s RNA editing function in mammals evolved as an exaptation driven by the emergence of *A3* and the arms race with retroviruses, while in birds, the lack of *A3* and lower ERV load resulted in *A1* inactivation and the absence of apoB RNA editing. Evidence of *A1*’s DNA editing activity in early mammals, such as opossums, and the correlation of *A3* expansion with the loss of *A1’s* antiviral function in primates support this hypothesis (Ikeda et al. 2017; Uriu et al. 2021; Moraes et al. 2022; Modenini et al. 2022). Hence, we expect the *A1* gene in birds to have undergone pseudogenisation, while the intact *A1* gene lacks functional motifs required for RNA targeting and experiences relaxed selection pressure. Considering DNA editing as the ancestral function, *A1*-specific mutations are expected in ERVs, whereas cis-acting factors associated with RNA editing machinery would be absent.

Here, we conduct a comprehensive in silico analysis of *A1* across birds, identifying and validating recurrent gene disruptions and catalytic inactivation in multiple lineages. We discover a correlation between *A1* presence/absence and the number of *A1*-associated DNA editing sites in the LTR retrotransposons of various bird species. Birds showing zero or minimal DNA editing sites exhibit signs of relaxed selection, suggesting a reduced evolutionary pressure from retroviruses.

## 2) Methods

### (a) Collection and recovery of *A1* sequences

To encompass avian diversity, we selected 81 species from 36 major bird orders (**Supplementary Table 1**) based on the availability of scaffold- or chromosome-level genomes, ensuring the representation of at least one species per order while avoiding closely related species based on divergence time. The divergence time for each species is obtained from the time-calibrated phylogenetic tree from Timetree (Kumar et al. 2022). Annotated *A1* and *A1*-like gene sequences were extracted from public databases such as Ensembl (Martin et al. 2023) and NCBI (Sayers et al. 2020) and validated against the respective reference genome using blastn.

A four-pass strategy was implemented with sequence similarity-based tools to recover the complete ORF of one-to-one orthologs of *A1* from all birds. The four-pass criteria include a query cover of 100%, complete ORF formation without premature termination codons (PTC), intact splice sites without exon boundary indels, and conserved synteny. Initially, blastn and blastx from NCBI BLAST+ (v2.9.0+) (Camacho et al. 2009) were used to identify the *A1* orthologs using the duck *A1* as a primary query against the bird genomes. Subsequently, the exon boundaries with intact splice sites were located using splice-aware aligners, namely exonerate (v2.2.0) (Slater and Birney 2005) and spaln (2.4.12) (Iwata and Gotoh 2012). In cases where the four-pass strategy could not recover an ORF with duck *A1* as the query, the process was iterated using a phylogenetically closer query derived from the time-calibrated phylogenetic tree. Moreover, the order and orientation of flanking genes are determined using the genomic position of *A1*’s BLAST hits, combined with annotation data from NCBI. For birds lacking annotation data, we performed TOGA (Kirilenko et al. 2023), which considers synteny as one of the features to distinguish orthologs from paralogs based on a reference annotation.

### (b) Identification of mutational events

To decipher the mutational events leading to the ORF-disrupting changes in all bird genomes lacking *A1*, we performed chromosome alignments using TOGA, with the duck genome as a reference. TOGA reports all indels (insertions and deletions) and substitutions located within the exons that disrupt the reading frame, as well as splice site mutations, but skips large repeats. To identify repeat insertions that overlap genic regions, RepeatMasker (v4.1.2) was used with the parameters -s (Slow search) and -species, specifying species-specific repeats (Tarailo-Graovac and Chen 2009). In addition to mutations that disrupt ORF, the integrity of catalytic and active sites was also examined. The identification of protein domains and catalytic sites was conducted through the batch CD-search interface, utilising the National Library of Medicine’s Conserved Domain Database (CDD) with default parameters (Wang et al. 2023).

### (c) Verification of gene-disrupting changes

To rule out the potential artefacts from assembly errors that could mimic gene-inactivating mutations, we screened unassembled raw short-read genomic datasets from multiple NCBI Short Read Archive (SRA) experiments (**Supplementary Table 2**) for each species using blastn. In contrast to SRA, long-read data can address large assembly gaps caused by repetitive elements and are less susceptible to GC bias. The assemblies of *Gallus gallus* and *Struthio camelus*, representing major clades with *A1* loss, and *Leptosomus discolor*, which lacks all *A1* exons, were assessed for long-range accuracy. Briefly, we mapped the long-read data from multiple PacBio and Nanopore datasets against the respective genomes using minimap2 (v2.16) (Li 2018) with default parameters. Subsequently, the resulting alignments were visually inspected using the Integrative Genomics Viewer (IGV) (Robinson et al. 2011). Reads longer than 10 kb were filtered, and the tiling path spanning the genomic region encompassing *A1* remnants and the adjacent genes was visualised using the UCSC Genome Browser (Nassar et al. 2023). Moreover, we utilised optical mapping data to examine structural variations in the genomic region of chicken containing *A1*, as it is not subject to sequencing bias due to PCR amplification and enzyme selection. Bionano optical maps for chicken were obtained from the GenomeArk database repository of the Vertebrate Genome Project. These maps were aligned to the genome using OMBlastMapper and visualised using OMViewer from OMTools (v1.4a) (Leung et al. 2017).

### (d) Evaluating the Transcriptional status of *A1*

To ascertain the loss of *A1*, we analysed RNA-seq data from various tissues of each species exhibiting the gene disruption. Initially, the transcriptional status of *A1* was assessed and manually examined using both tissue-specific and aggregated RNA-seq exon coverage tracks, along with intron features and intron-spanning reads available in the genomic tracks at NCBI. For species lacking tissue-specific tracks, transcriptomic datasets were retrieved from the NCBI SRA database and aligned to their respective genomes using the STAR (version 2.7.0d) mapper (Dobin et al. 2013). The per-base coverage of resultant alignments was plotted using the ggplot2 R package (Wickham 2016), and the splicing junctions were visually inspected using IGV.

### (e) Clustering and phylogenetic analysis of *A1* homologs

To investigate the relationship of *A1*-like within the *AID*/*APOBEC* family, we performed clustering using amino acid sequences from *A1* homologs across vertebrates, including 106 Birds (53 *A1*, 53 *A1*-like), 46 Mammals, 40 Testudines, 6 Crocodylia, 13 Squamata, 2 Amphibians and 3 Lungfish. Other *AID*/*APOBEC* members, such as *AID* (36), *A2* (60), *A3* (36) and *A4* (30), were also included in the analysis as outgroups. In total, 379 full-length CDS were retrieved from NCBI, Ensembl, and OrthoDB (Tegenfeldt et al. 2025) and utilised for analysis after quality control checks, including assessments for genome gaps, exonic repeats, synteny, and RNA expression, to ensure correct orthology and avoid annotation errors. Clustering was performed using CLANS, which performs all-against-all pairwise BLAST searches and visualises the matches as a similarity network (Frickey and Lupas 2004). To further examine the origin and evolution of the *A1*-like gene, we conducted a phylogenetic analysis using CDS. Initially, all CDS sequences were aligned using Guidance (V2.02) with MAFFT as the alignment tool, employing 1000 bootstrap iterations (Sela et al. 2015). A maximum likelihood tree was then constructed using IQ-TREE (V2.3.2) with automated extended model selection (MFP) and 5000 ultrafast bootstrap replicates (Minh et al. 2020). The final tree was visualised, collapsed, and labelled using Figtree (V1.4.4) (Rambaut et al. 2018).

### (f) Identification of *A1*-associated DNA editing

To test whether *A1* acts as a DNA editor in birds, we examined bird genomes for G-to-A mutations, specifically focusing on long terminal repeats (LTRs). *A1* is known to target LTR retrotransposons by editing “C” to “U” in single-stranded antisense DNA during reverse transcription, resulting in “G” to “A” changes upon genomic reintegration. To detect these DNA edit sites, we adapted a mismatch-based method from Knisbacher & Levanon (Knisbacher and Levanon 2016). For an LTR sequence to be considered edited, it must have an unedited ancestral element in the genome with high sequence similarity, except for G-to-A mismatches at editing sites. We used blastn to generate pairwise alignments among closely related LTRs within the same subfamily, setting parameters to an E-value of 1E-50, strand to plus, dust to no, soft_masking to false, and num_alignments to 250. Alignments with clusters of at least five consecutive G-to-A mismatches, referred to as “clusters”, were identified, as *APOBEC* mutations typically occur in clusters rather than being uniformly distributed across the target sequence. Additional filters were applied to identify precise sites that are strongly associated with *APOBEC* activity. To ensure the accuracy of the detected edit sites, three negative controls were used: the number of C-to-T edit sites in bird LTRs, G-to-A edit sites in bird DNA transposons and invertebrate LTRs. To normalise the number of GA edit sites in each bird, we applied a log transformation to reduce skew, normalising by both CT sites and total LTR size, as defined by the equation,

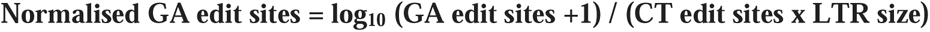

To account for species with zero GA edit sites, we added +1 to ensure their inclusion in the log transformation.

## 3) Results

### (a) *APOBEC1* in birds do not edit RNA

#### (1) Birds lack sites in *A1* required for RNA editing

To investigate the evolutionary divergence of *APOBEC1* (*A1*), we collected and compared the coding sequences of synteny-confirmed one-to-one orthologs from birds and mammals, the latter being the only lineage in which *A1* function has been experimentally established. Notably, all avian *A1* orthologs lack the C-terminal segment encompassing approximately 38–43 amino acids that correspond to the A1HD domain (**Fig. 1**). In mammals, this domain forms a unique hydrophobic fold, specific to *A1*, and is essential for dimerisation. This region contains several residues critical for RNA editing, including R198, Q200, R207, and H214. Similarly, the residues L203, L208, and L210 in the same region constitute a conserved leucine-rich motif (**Supplementary Table 3**). Additional residues, such as R197, contribute to structural stability, while L218 is required for RNA binding. Although this region is dispensable for editing activity, monomeric enzyme remains active, often exceeding the dimer in catalytic efficiency (Wolfe et al. 2020). However, the lack of a cofactor binding site removes the high target specificity required for RNA editing, and the inability to form a dimer reduces the availability of *A1*, making it more prone to degradation.

**Figure 1:**
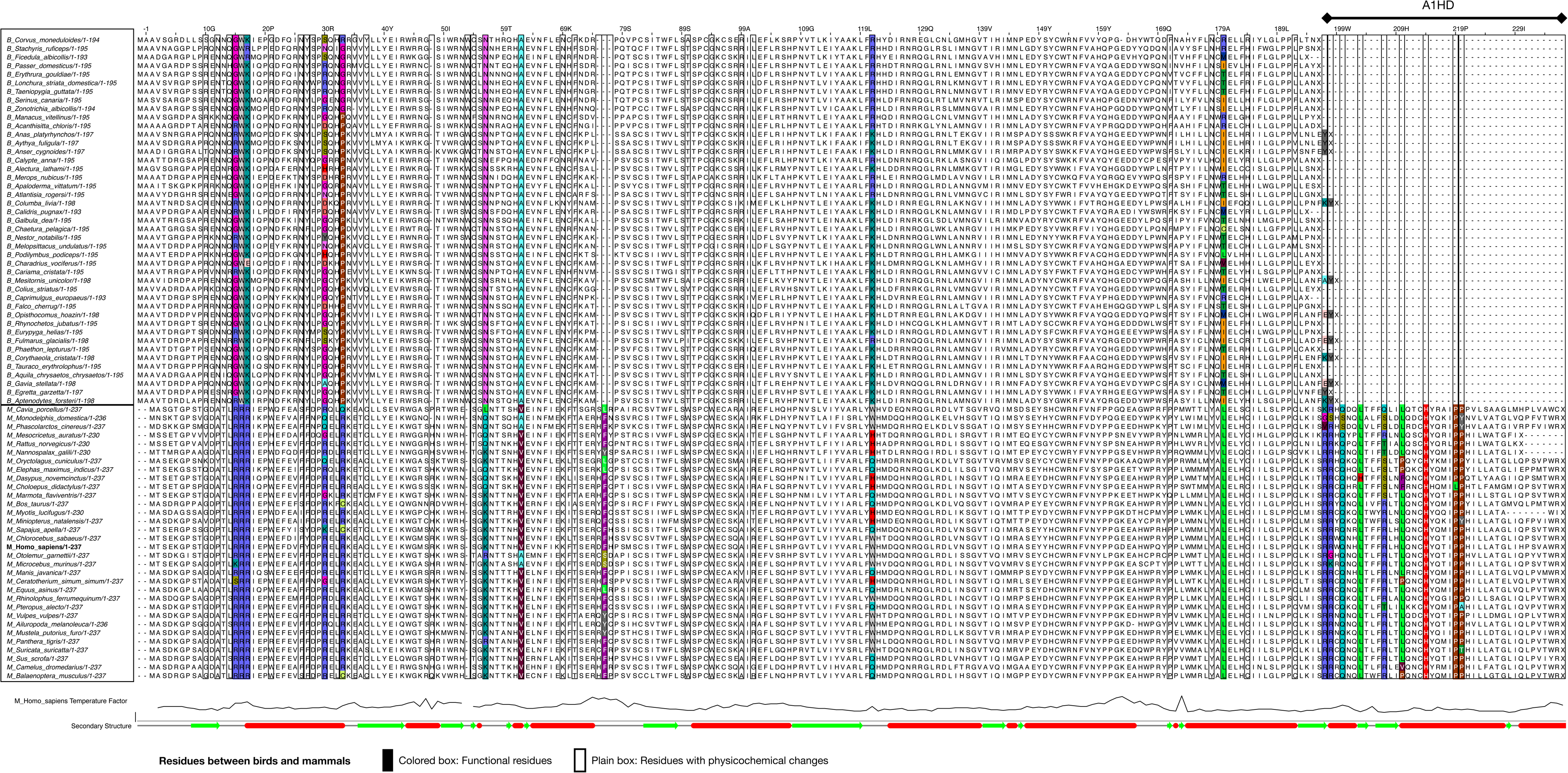
Absence of the C-terminal domain and functionally relevant sites of *A1* in birds. A multiple sequence alignment of the *A1* protein from birds and mammals with intact *A1* is presented here. Amino acid differences between mammals and birds are represented by colored boxes marking functionally important residues identified from previous studies, while uncoloured boxes indicate substitutions involving physicochemical changes in charge, hydrophobicity, volume, or chemical nature. Residue positions are based on the human *A1* protein, with functional sites and regions relevant to apoB RNA editing. The C-terminal region, absent in all bird sequences compared to mammals, contains critical sites responsible for RNA editing and A1CF cofactor binding (R198, Q200, R207), dimerisation (residues 196–236), and structural stability (R197). This region forms the A1HD domain and includes a conserved leucine-rich region that overlaps with the nuclear export signal. Additional functional sites outside the A1HD domain, important for RNA editing (R15, R16, R17) and RNA substrate selection (F76/Y76, W121/H121/Q121), are also labelled. These collective disruptions suggest that even in birds with intact *A1*, RNA editing, dimerisation, and cofactor interactions are likely absent.

Secondly, several conserved residues upstream of the A1HD domain in mammalian *A1*, which are essential for RNA binding and editing, are absent in avian *A1*. These include R15, R16, R17, K56, V62, and L180—residues known to be critical for RNA editing in mammals. The basic residues R15, R16, and R17, at least two of which are required for RNA binding, editing, and A1CF interaction, are highly conserved across mammals but absent in all bird sequences. Notably, this tripeptide (R15–R17) also functions as a nuclear localisation signal (NLS). Furthermore, R30, a residue contributing to structural stability like R197, is missing in all avian *A1* sequences except for five passerines, which have high LTR DNA edit sites (see section b of the results). Another key difference is at position 121, where mammals possess W121—a residue located in loop 7 and critical for RNA editing but not DNA deamination. While mammals tolerate substitutions of W with H or Q at this site, birds exhibit R121 or K121. Recent evidence suggests that this position plays a role in determining substrate preference between RNA and DNA, indicating that avian *A1* likely functions as a DNA editor rather than an RNA editor (Wolfe et al. 2020).

Finally, we identified several amino acid substitutions not previously linked to specific functions but likely to influence *A1* structure and activity through alterations in physicochemical properties, including charge, hydrophobicity, volume, and chemical nature (**Fig. 1**, **Supplementary Table 4**). To assess their potential functional impact, we performed PROVEAN analysis using human *A1* as the reference sequence. This revealed a higher proportion of residues in avian *A1* predicted to be deleterious compared to mammalian *A1* (**Supplementary Table 5**). Collectively, these findings highlight marked functional divergence between avian and mammalian *A1*, with avian proteins specifically lacking the features required for the precise RNA binding and editing characteristic of their mammalian counterparts.

#### (2) Lack of cis-acting elements in target RNA

To assess the potential for apoB RNA editing in birds, we compared apoB RNA sequences from birds and mammals, as conserved cis-acting elements within apoB are known to be critical for RNA editing. The sequence alignment revealed multiple nucleotide substitutions in the mooring sequence, efficiency elements, and A-rich regions of avian apoB transcripts, including the absence of the target cytidine (**Supplementary Figure S1**). Importantly, while the mooring sequence, which serves as the recognition site for the target base, is highly conserved in mammals, birds exhibit several nucleotide variations, suggesting that apoB RNA editing is likely absent in all birds.

### (b) *A1* as a DNA editor in birds

To assess whether *A1* functions as a DNA editor, we searched for *A1*-associated mutations in bird genomes, focusing on LTR retrotransposons, as *A1* has previously been shown to edit these elements (Knisbacher and Levanon 2016; Ikeda et al. 2017). We initially screened for G-to-A mutations within LTRs and applied multiple validation filters to ensure that the detected mutations were attributable to *APOBEC*-mediated activity. We found that the number of DNA editing sites was higher in certain passeriform species compared to other avian orders, independent of total LTR content, with a maximum of 46,042 edit sites identified in Zebra finch (*Taeniopygia guttata*) (**Supplementary Table 6**). In contrast, 73 out of 81 bird species exhibited fewer than 1,000 sites, including 43 species with no detectable edit sites. Notably, species with a higher endogenous retrovirus (ERV) content also showed greater numbers of edit sites, suggesting a link between *A1* activity and retroviral repertoire variation in birds (**Supplementary Table 7, Supplementary Figure S2**). As expected, all 16 invertebrate species used as negative controls exhibited no DNA editing (**Supplementary Table 8**). In contrast, non-mammalian vertebrates such as alligator (*Alligator mississippiensis*) (22,417 sites) and anolis lizard (*Anolis carolinensis*) (11,872 sites) displayed substantial DNA editing activity. Mammals also showed high numbers of edit sites, likely reflecting the combined contributions of both *A1* and *A3* enzymes. Taken together, these results suggest that DNA editing represents the ancestral function of *A1*, with distinct lineages—including birds, squamates, crocodilians, and mammals—retaining DNA editing signatures, whereas RNA editing may have evolved secondarily as an exaptation specific to mammals.

### (c) Inactivation of *A1* in Birds

#### (1) Inactivation in Paleognathae and Galliformes

Given that *A1* likely functions as a DNA editor in birds, providing defence against retroviruses and retrotransposons, the observed association between ERV load and the extent of DNA editing suggests that changes in retroviral repertoire may have influenced the evolutionary pressures acting on *A1*. Specifically, bird species with a lower ERV burden may have experienced relaxed selection on *A1*, potentially leading to gene loss, whereas species with a higher ERV load may have been subject to purifying (negative) selection to maintain *A1* function. To investigate the evolution of *A1* in birds, we extracted and inspected the coding region from 81 representative birds. The four-pass strategy revealed *A1* gene loss in two major groups of birds, namely Galliformes and Palaeognathae, which include multiple orders. In ducks, *A1* comprises five exons, with exon 3 spanning approximately 66% of the gene, exon 4 comprising 20%, and exons 1, 2, and 5 collectively making up the remaining 14%. All six species representing Palaeognathae shared the deletion of the first three exons of *A1*, while Greater rhea (*Rhea americana*) and White-throated tinamou (*Tinamus guttatus*) have additionally lost exon 4 and exon 5, respectively (**Fig. 2**). Retained exons in these lineages also harbour frameshifting indels and in-frame stop codons, with some mutations shared across species and others remaining exclusive to a single lineage.

**Figure 2:**
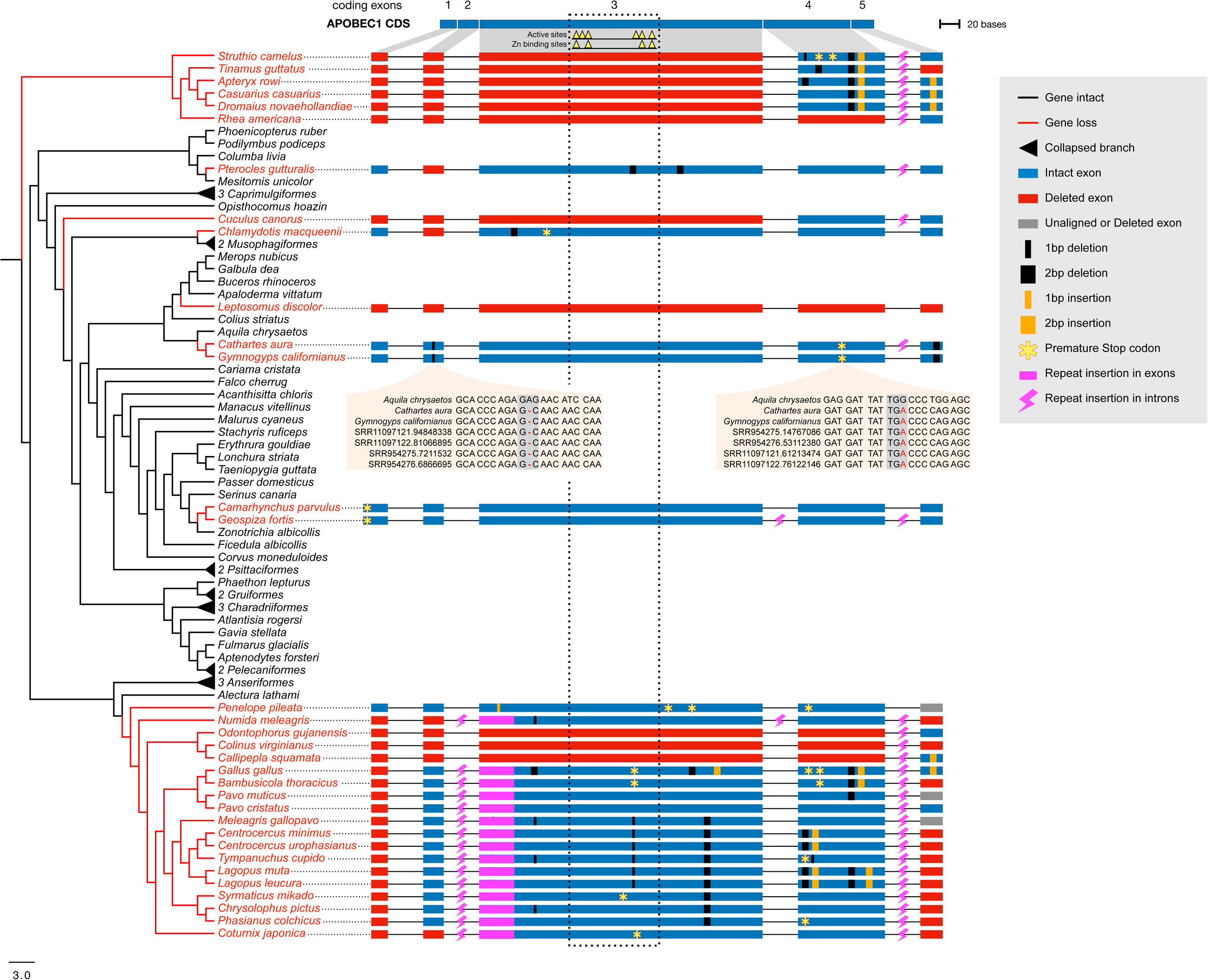
Lineage-Specific and Independent Loss of *A1* in Birds. The phylogeny of all birds analysed in this study (left) is shown, along with a schematic representation of *A1* exons and introns (intron lengths not to scale). At the top, a coding sequence (CDS) bar highlights active and zinc-binding sites within *A1*. Red colored branches and labels represent birds with *A1* loss, while those with intact *A1* are marked in black. ORF-disrupting mutations are colour-coded: red bars indicate whole exon deletions, blue bars indicate intact exons, black marks indicate deletions, and yellow indicates insertions, with indel length proportional to actual size. Premature termination codons (PTC) are denoted by yellow stars outlined in red and are distinct from PTC caused by indels. Mutations within active and zinc-binding sites are enclosed in rectangular boxes. Repeat insertions are marked in magenta, with exon-overlapping repeats represented as bars and intron-overlapping repeats as thunderbolt symbols. This figure illustrates the inactivation of *A1* in birds through the accumulation of shared and independent mutations, with sufficient gene remnants remaining to facilitate accurate identification.

In Galliformes, 19 species, with the exception of Australian brushturkey (*Alectura lathami*), exhibit *A1* disruptions, suggesting that inactivating mutations occurred before the divergence of the common ancestor of white-crested guan (*Penelope pileate*) and Australian brushturkey (*Alectura lathami*) (**Fig. 2)**. Importantly, no mutations are shared between Palaeognathae and Galliformes, indicating parallel inactivation in the ancestral lineages of each group. All 19 Galliformes species exhibit a shared deletion of exon 1, while Odontophoridae, represented by three species, also show deletions of exons 2, 3, and 4. Additionally, repeat insertions were observed in exon 3 in ten species, and repeats in the second and fourth introns are shared across all Galliformes species. Numerous in-frame stop codons and short indels (1–2 bp) were identified in exons 3 and 4, with most mutations shared by at least two species. Despite an extensive search of the genome and both short- and long-read datasets, no additional translocated copies of *A1* were found in the chicken.

To rule out the possibility of assembly errors or SNPs being misinterpreted as inactivating mutations, we screened raw SRA reads from both groups. All inactivating mutations were supported by reads coming from at least two different SRA datasets (**Supplementary Table 2**). Furthermore, long-read data (PacBio and Nanopore) from South African ostrich (*Struthio camelus australis*) and chicken (*Gallus gallus*), representing each group, were aligned against the respective genomes and showed no drop in coverage across *A1* and flanking genes (**Supplementary Figure S3**). Nanopore reads mapped to an approximately 45 kb region of chicken (*Gallus gallus*), spanning the genes *A1*, along with upstream *NANOG* and downstream *AICDA*, showed high coverage (**Supplementary Figure S4**). Similarly, the more accurate PacBio reads also displayed consistent coverage across the *A1* region and adjacent genes (**Supplementary Figure S5**). Additionally, chicken optical maps displayed no structural variations across the genomic region encompassing *A1* (**Supplementary Figure S6**). Taken together, this multi-platform evidence strongly supports the loss of *A1* in these avian groups.

#### (2) Independent loss of *A1* in birds

We have also found inactivating mutations in eight bird species, including two Passeriformes (*Camarhynchus parvulus, Geospiza fortis*), two Accipitriformes (*Cathartes aura, Gymnogyps californianus*), *Pterocles gutturalis* (Ciconiiformes), *Leptosomus discolor* (Coraciiformes), *Cuculus canorus* (Cuculiformes) and *Chlamydotis macqueenii* (Gruiformes). Importantly, all these species occupy different branches of the phylogeny and lack shared mutations, indicating independent losses in their common ancestors. In the genome of *Leptosomus discolour*, both short-read and PacBio HiFi reads validate the ∼8.2 kb deletion that removes all five exons of *A1* reads (**Supplementary Figure S7**). Using *A1* from closely related species as a query showed no evidence for deleted exons in unassembled reads, while mapped long reads revealed no gaps in the assembly covering the flanking gene *AICDA*. Other species with exon deletions include *Cuculus canorus*, which has deletions of exons 1, 2 and 3, whereas both *Pterocles gutturalis* and *Chlamydotis macqueenii* exhibit deletion of exon 2. We also identified species-specific indels and in-frame stop codons in retained exons, all of which are located within the first 50% of the entire coding sequence. In total, we found 33 individual bird species with *A1* disruptions, comprising six from Palaeognathae, 19 from Galliformes, and eight independent losses from various lineages.

#### (3) Disruption of catalytic sites in bird *A1*

To evaluate the status of functional domains and catalytic sites in the *A1* gene, we employed NCBI’s batch CD-search. For species with an intact *A1*, amino acid sequences were analysed, whereas for bird species with *A1* disruptions, translated coding DNA sequences in all three reading frames were examined to rule out the possibility of intact catalytic site formation in any frame. The hits from CD-search results revealed that bird species with an intact *A1* exhibit specific hits corresponding to *APOBEC4*-like *AID*/*APOBEC*-like deaminases (PF18774). The catalytic sites HxE and PCx2-4C, essential for deamination, were found to be intact in the third exon across all 48 species with an intact *A1*, whereas CD-search failed to detect them in 29 bird species with *A1* disruptions. Among the 33 species with loss, three species lacked the *A1* gene entirely, while 11 species, including all Palaeognathae, three Galliformes, and two species from independent lineages, lacked exon 3, which contains the catalytic sites (**Fig. 2**). For the remaining 22 bird species, all three reading frames of *A1* remnants were manually inspected, revealing no intact catalytic motifs in any frame, except in four species (two Passeriformes and two Accipitriformes). Despite retaining intact catalytic sites in one of the three reading frames, these species exhibited inactivating mutations upstream of the catalytic sites, likely causing a frameshift and rendering the enzyme inactive. These findings suggest a loss of enzymatic activity in bird species with *A1* disruptions, further supporting the inactivation of this protein and providing insights into its functional implications.

#### (4) Synteny and transcriptional status of *A1*

As remnants of a disrupted gene with conserved synteny can serve as a strong indicator of gene loss, we examined the order and orientation of *A1* and its flanking genes. Syntenic analysis revealed a conserved gene order and orientation, with *SLC2A3*, two *NANOG* and *A1*-like positioned upstream and *AICDA*, *MFAP5* and *RIMKLB* located downstream to *A1* in all bird species except Galliformes, which lack *A1*-like (**Fig. 3**). Species exhibiting a complete loss of exons, such as *Leptosomus discolor* and *Colinus viginianus*, also retained this conserved gene order, with chain alignment extending from *AICDA* to the downstream intergenic region of *A1* (**Supplementary Figure S8**).

**Figure 3:**
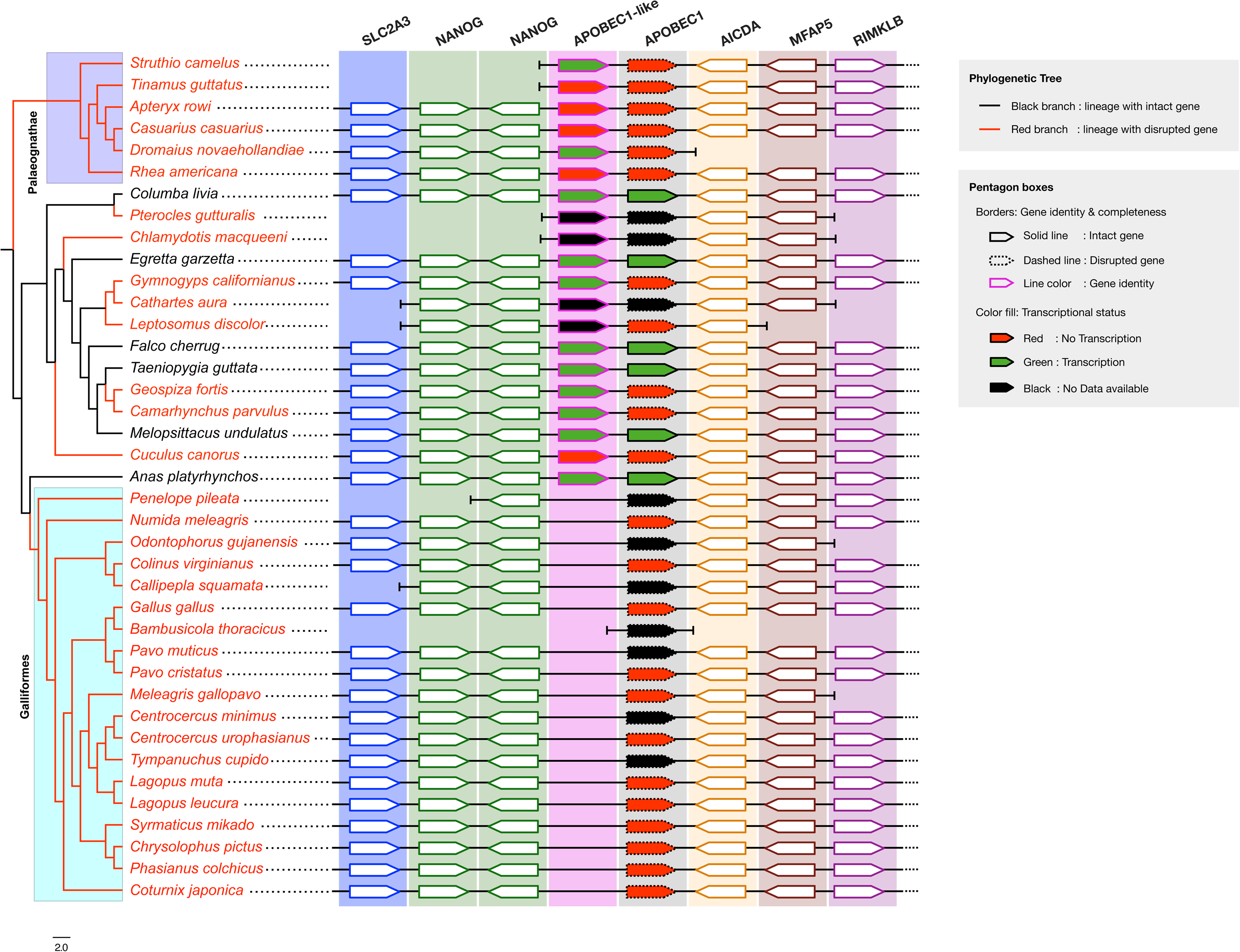
Synteny and Transcriptional Status of *A1* in Birds. On the left, a phylogenetic tree illustrates the loss of the *A1* gene in birds, marked in red, along with the closest species that retain an intact *A1*. On the right, each pentagon represents a gene, with the border colour indicating gene identity and the border style (solid or dashed) representing gene completeness. Dashed borders indicate open reading frame (ORF) disruptions, while solid borders indicate complete genes. Gene orientation is shown by the direction of the pentagon’s apex, with right-pointing apexes representing a 5’ to 3’ direction and left-pointing apexes representing a 3’ to 5’ direction. Conserved synteny is observed, with *A1*-like, *NANOG* (two copies), and *SLC2A3* located upstream of *A1*, and *AICDA*, *MFAP5*, and *RIMKLB* downstream, confirming these *A1* genes as one-to-one orthologs. Notably, *A1*-like is found in all birds except Galliformes. RNA expression is visualised by box colour, where green indicates active transcription, red indicates no transcription, and blue represents unavailable RNA data. All bird species with *A1* loss show no RNA transcription across tissues analysed, while the closest species with intact *A1* displays strong transcription across multiple tissues, underscoring the inactivity of the lost *A1* gene.

To rule out the possibility of *A1* being expressed despite inactivation mutations, we assessed RNA-seq exon coverage across all available tissues using the NCBI Genome Viewer and IGV. Under normal conditions, genes containing PTCs are targeted by nonsense-mediated mRNA decay (NMD), a process that degrades mRNAs, preventing the formation of truncated, dysfunctional proteins (Chang et al. 2007). Although some transcripts can evade NMD and undergo translation, a PTC located at least 50–55 nucleotides upstream of the last exon-exon junction is generally sufficient to trigger NMD (Hwang and Kim 2013). Here, we identified multiple bird species with PTCs in *A1*, occurring on or before the third exon—well within the effective range for NMD, at least 116 nucleotides upstream of the final intron. Additionally, some bird species showed the loss of one or more entire exons, as exon skipping is another natural NMD target [80]. As expected, our screening indicated a complete absence of RNA expression in all 33 bird species with *A1* disruptions, whereas the phylogenetically closest species with intact *A1* demonstrated robust RNA expression across multiple tissues (**Fig. 3**). Interestingly, *Alectura lathami*, the only Galliform species with an intact *A1*, also exhibited *A1* expression in blood, suggesting the loss event occurred after its divergence from other Galliformes. Taken together, these findings indicate that conserved synteny confirms that the lost gene is a one-to-one ortholog of *A1* in bird species, while the absence of RNA expression across various tissues indicates the inactivity of the lost gene.

#### (5) *A1* loss is correlated with DNA edit sites

As birds display varying levels of APOBEC-associated DNA editing, we examined whether these patterns correlate with the presence or absence of the *A1* gene. A phylogenetic correlation analysis was conducted to examine the relationship between *A1* status and the number of GA editing sites. To account for interspecies variation and reduce skew, the number of GA sites was log-transformed and normalised by both the CT sites and the total LTR sequence length. Following normalisation, the number of bird species suitable for phylogenetic regression decreased from 81 to 31 due to the zero CT edit sites in several species. The regression analysis revealed a positive and statistically significant correlation between *A1* presence and GA editing abundance (*p* < 0.05, *n* = 31). To test the robustness of this relationship, two independent methods, namely MPLE (*p* = 0.0151) and IG10 (*p* = 0.0156), were applied, both yielding consistent results. Collectively, these findings indicate that the loss of *A1* is functionally linked to a reduction in DNA editing and support its role as a DNA editor in birds.

### (d) *A1*-like genes in birds

Genomic annotations of several bird species identified an *A1*-like gene upstream of *A1*, between *NANOG* and *A1*, alongside six conserved syntenic genes. Genomic annotations for this gene vary widely across bird species, with some lacking it entirely, while others classify it as *A1*-like, *A1*, a non-coding RNA, or a merged *A1* and *A1*-like gene with a double ZDD domain (**Supplementary Table 9**). A homology-based search confirmed its presence in all bird species examined, except Galliformes, with *Alectura lathami* as the only exception retaining both *A1* and *A1*-like. RNA expression data further distinguishes *A1*-like, as it is transcribed in multiple tissues, while *A1* expression remains undetected, including in species that have undergone *A1* loss (**Supplementary Table 9**). Additionally, in species annotated with a double ZDD domain, the intron feature track shows RNA-seq reads supporting splice junctions mapped to all introns except one, corresponding to the intergenic region between *A1* and *A1*-like (**Supplementary Figure S10**). Collectively, these findings suggest that *A1*-like represents a distinct gene from *A1*, broadly distributed among bird species.

To investigate the evolutionary relationship between *A1*-like, *A1*, and other *AID*/*APOBEC* family members, we performed a clustering analysis of *A1* homologs across vertebrates, using *AID*, *A2*, *A3*, and *A4* as outgroups. Since *A1* was first identified and characterised in mammals, we adopted a nomenclature to distinguish *A1* from *A1*-like across tetrapods, referring to “*A1*” as the single-copy gene in mammals and the downstream copy in birds (bird *A1*), “*A1*-like” as the upstream copy in birds (bird *A1*-like), and *A1* homolog for all copies in other tetrapods (e.g., Crocodylian *A1* homolog) (**Supplementary Table 10-11**). Clustering analysis revealed that *A1* homologs from all tetrapod groups, including bird *A1*-like, formed a distinct supercluster separate from *AID*, *A2*, *A3*, and *A4*, with bird *A1*-like clustering separately from bird *A1*, confirming its identity as a distinct *A1* paralog (**Fig. 4a**). Additionally, Crocodylian and Testudine *A1* homologs formed an intermediate cluster between bird *A1* and *A1*-like, with some sequences clustering closer to *A1* and others to *A1*-like. These findings indicate that the upstream gene adjacent to bird *A1* represents a true *A1* paralog and that *A1*-like divergence is specific to Archelosauria.

**Figure 4:**
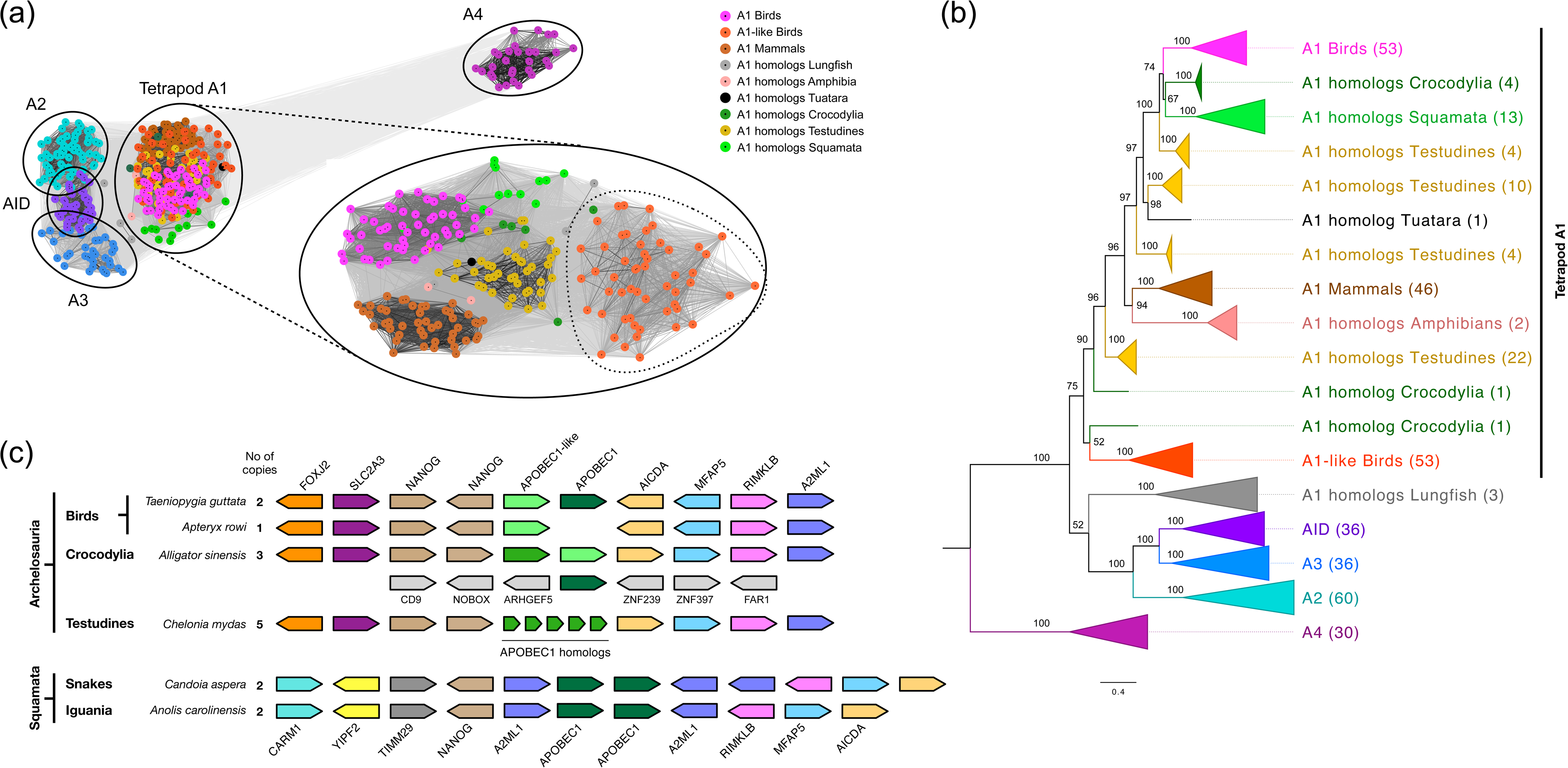
*A1*-like in Birds. **(A)** Clustering analysis of the *AID*/*APOBEC* family reveals that major family members form distinct clusters, with each dot representing a protein coloured based on its family affiliation. The zoomed-in portion highlights the *A1* cluster, which contains subclusters of *A1* homologs across vertebrates, coloured by taxonomic group. Notably, bird *A1*-like (magenta) clusters separately from bird *A1* (orange), suggesting the formation of a distinct paralog within birds. Similarly, crocodilian and Testudine *A1* homologs exhibit divergence, with some clustering closer to bird *A1* and others showing bird *A1*-like characteristics, indicating that the *A1*-like divergence is specific to Archosauria. **(B)** The phylogenetic tree of the *AID*/*APOBEC* family was constructed using coding sequences (CDS) and coloured by family members and taxonomic groups. Bootstrap values are provided at each node, with the number of sequences indicated in parentheses. The tree shows all *A1* homologs branching separately from other family members, except for the lungfish *A1* homolog. The *A1* lineage exhibits monophyly, with bird *A1* and *A1*-like forming a polyphyletic branching pattern. *A1* homologs in non-avian tetrapods exhibit multiple distinct branches, indicating paralogous divergence and the dynamic evolution of the *A1* gene. **(C)** Generalised synteny analysis of major amniote groups, illustrating gene order conservation between Archelosauria and Squamata. *A1*-like is found exclusively in Archelosauria, further supporting its lineage-specific divergence.

To further investigate the origin and evolution of *A1*-like, we constructed a phylogenetic tree. *A1* homologs were found to form a monophyletic clade separate from other *AID*/*APOBEC* family members, with the exception of the lungfish *A1* homolog, which branched as a sister group to *AID* (**Fig. 4b**). This pattern is consistent with previous findings that *A1* arose through duplication of *AID*. Among tetrapods, only Crocodylian and Testudine *A1* homologs formed close phylogenetic relationships with bird *A1*-like, while other tetrapod *A1* homologs grouped with bird *A1*, suggesting that *A1*-like divergence is restricted to Archelosauria. Additionally, synteny analysis revealed high copy number variation in *A1* within Archelosauria, ranging from a single copy in some birds to six copies in Testudines. Birds and Testudines retained a conserved syntenic block, whereas Crocodilians had one copy at a different genomic locus (**Fig. 4c**). Similarly, squamates possess two *A1* copies, but with a different gene order, featuring *A2ML* as flanking genes and an inverted arrangement of three downstream *A1* genes. In contrast, mammals and amphibians possess a single *A1* copy with a distinct syntenic organisation, suggesting *A1* duplication might have occurred in the amniote ancestor after divergence from amphibians, followed by lineage-specific rearrangements. Notably, *A1* homologs from Crocodylian formed three branches, with two forming direct sister groups to bird *A1* and bird *A1*-like, respectively. In contrast, Testudines *A1* copies split into four polyphyletic clades, showing varying degrees of relatedness to bird *A1* but lacking a direct sister-group relationship with birds (**Fig. 4b**). Taken together, these findings highlight the dynamic evolution of *A1*, where paralogs likely diversified independently following duplication, resulting in distinct clades, copy number variation reflecting gene loss and amplification, disrupted synteny from genomic rearrangements, and polyphyletic branching shaped by differential selection pressures.

### (e) *A1* loss is not linked to *ApoE* loss

To determine whether *A1* loss in birds is evolutionarily linked to *ApoE* loss, we searched avian genomes for *ApoE*, as *A1*-edited apoB48 lipoproteins are cleared from circulation via *ApoE*. *ApoE* was identified in multiple bird species, including Palaeognathae and *Gallus gallus*, both of which lack the *A1* gene. In *Gallus gallus*, *ApoE* was missing in the reference genome (GCF_016699485.2) but was successfully recovered from the white leghorn layer genome (GCA_016700215.2). Since missing genes are often GC-rich and enriched in tandem repeats, we analysed apoE GC content across tetrapods and found that birds exhibit extreme GC richness, with all sequences exceeding 70%, which likely explains their absence in certain genome assemblies (**Supplementary Figure S11**). Although *A1* synteny in birds showed partial conservation with crocodilians and mammals, apoE scaffolds in birds were fragmented, reflecting sequencing challenges in GC-rich regions. These findings indicate that *A1* loss does not correlate with apoE loss, suggesting that *A1* in birds did not evolve to do apoB RNA editing.

## 4) Discussion

In this study, we report the lineage-specific and independent inactivation of the *A1* gene across all bird orders. Previous studies have reported the apparent absence of the *A1* gene among birds, stemming from the inability to detect *A1* in chicken enterocytes (Teng and Davidson 1992). Subsequent investigations identified *A1* in several mammalian species, suggesting that it may be a mammalian-specific gene. However, recent studies challenge the earlier notion that *A1* is a mammalian-specific gene, revealing its presence in reptiles, birds, turtles, amphibians, and lungfish. Some species exhibit amplified *A1* paralogs, while others, particularly certain amphibians and birds, display gene loss (Conticello et al. 2005; Krishnan et al. 2018). In birds, evidence suggests *A1* may have pseudogenised in certain lineages, but it remains unclear which birds have lost *A1* and what mechanisms drove its inactivation. Despite discoveries since 1990 (Tarugi et al. 1990), including *A1*’s absence in chickens and presence in *Taeniopygia guttata* (Severi et al. 2011), no comprehensive study has yet mapped the full evolutionary dynamics of *A1* in bird genomes. This is the first study to establish the inactivation mutations that lead to the loss of *A1* throughout all bird orders.

In total, we identified 33 bird species with *A1* disruption, comprising six from Palaeognathae, 19 from Galliformes, and eight independent losses from various lineages. We provide evidence for inactivation from the genome, unassembled SRA data, long reads, and optical maps, ruling out the possibility of inactivation due to polymorphism, GC bias, sequencing and assembly errors. The *A1* coding sequences in Palaeognathae and Galliformes reveal at least two shared mutations, including exon deletions and repeat insertions. This observation aligns with phylogenetic hypotheses suggesting two initial inactivation events followed by the random accumulation of additional mutations. A gene is considered lost when it has acquired multiple gene-inactivating mutations and has a lineage descending from an ancestor with an intact gene (Sharma et al. 2018). However, in addition to the physical disruption, we also demonstrate the possibility of a loss of function. Despite the *A1* targets, the disruption of catalytic and Zn-binding sites essential for the deamination reaction implies a functional loss, even if a truncated or variant of *A1* is somehow expressed.

Inactivated gene remnants with conserved synteny offer strong evidence for the loss of one-to-one orthologs. In this study, we show that the gene order and orientation of both upstream and downstream genes to *A1* are highly conserved across birds. To further validate *A1* loss, we analysed available RNA-seq data, indicating the absence of alternative splicing patterns. Under normal conditions, genes containing PTCs are targeted by NMD, a process that degrades mRNAs, thereby preventing the formation of truncated, dominant-negative, or dysfunctional proteins (Chang et al. 2007). Although some transcripts can evade NMD and undergo translation, a PTC of at least 50–55 nucleotides upstream of the last exon-exon junction is generally sufficient to trigger NMD. Here, we identified multiple bird species with PTCs in *A1* occurring on or before the third exon, well within the effective range for NMD, at least 116 nucleotides upstream of the final intron. Additionally, some birds show loss of one or more entire exons, as exon skipping is another natural NMD target. As expected, all 33 bird species with *A1* loss showed no RNA expression across multiple available tissues, whereas the closest species with intact *A1* displayed clear RNA expression.

In addition to *A1*, we identified an *A1*-like gene upstream of *A1* in all examined bird species, with the exception of the Galliformes. Among Galliformes, only *Alectura lathami* retains both *A1* and *A1*-like. Although experimental validation of *A1*-like is currently lacking, RNA-seq data with exonic and intron-spanning reads indicate that *A1*-like is distinct from *A1*. Despite inconsistencies in annotation, clustering and phylogenetic analyses consistently showed that *A1*-like is most closely related to *A1* rather than to *AID*, *A2*, *A3*, *A4*, or *A5*, supporting its identity as an *A1*-like gene. The close genomic location of *A1*-like adjacent to *A1* and the presence of *A1* in early amniotes, while *A1-*like appears to be restricted to the Archelosauria lineage, suggest that *A1*-like likely arose from an ancestral duplication of *A1*. Within Archelosauria, the copy number and sequence of *A1*-like vary greatly, with Testudines possessing up to six copies and Galliformes entirely lacking *A1*-like. In birds, *A1*-like is expressed across various tissues, although its specific function remains unclear. Together, these findings indicate a dynamic evolutionary history for *A1*, encompassing duplication, the emergence of a distinct *A1*-like gene, and recurrent gene losses across lineages.

The physiological function of *A1* is well established in mammals, where it catalyses the deamination of C-to-U in apoB RNA, creating a stop codon leading to a truncated protein called apoB48. *A1* is highly conserved in placental mammals, as apoB48 is a key player in lipid transport, and several mutations in the gene have been shown to lead to lipid disorders (Lo and Coschigano 2020). However, in birds such as chickens, both *A1* and apoB RNA editing are absent despite the presence of editing cofactors *A1CF* and *RBM47*. In contrast to chicken, intact *A1* is retained in *Taeniopygia guttata* while lacking apoB RNA editing, suggesting an alternate function might exist for *A1* in birds. Here, we show that several cis-acting regions in the apoB RNA of all birds exhibit several nucleotide differences relative to mammals. Importantly, the mooring sequence, which acts as a recognition site for the target base, is highly conserved in mammals as virtually all mutations in this region are shown to abolish apoB RNA editing. All birds exhibit several nucleotide changes in the mooring sequence, including birds with intact *A1*, suggesting apoB RNA binding and editing are likely absent even in the presence of functional *A1*. Additionally, apoE, an apolipoprotein that acts as a ligand for the uptake of lipids from apoB48-containing lipid particles, is also thought to be absent in birds (Liu et al. 2019). The lack of apoE in birds was first reported in pigeons in 1985 (Barakat and St. Clair 1985) and later confirmed in chicken (*Gallus gallus*) and turkey (*Meleagris gallopavo*) using in silico methods (Liu et al. 2019). While these studies indicate the absence of apoE, genomic annotation reveals the presence of apoE in a few birds, primarily from the Palaeognathae. Our findings show that some birds with intact *A1* lack the apoE gene. If birds possess apoB RNA editing by *A1*, the lack of apoE will cause lipid disorders, as apoE is the only ligand that can uptake apoB48 particles. Thus, having *A1* in the absence of apoE is more detrimental than lacking *A1*, suggesting that apoB editing by *A1* is likely absent in birds.

We observed *APOBEC*-associated DNA editing signatures within ERV regions across several bird species, Crocodylia (*Alligator mississippiensis*), and Squamata (*Anolis carolinensis*), suggesting an ancestral role of *A1* as a DNA editor. Among the only APOBEC family members present in birds—*AID, A2,* and *A1—A2* lacks known deaminase activity, whereas *AID* is established to deaminate only immunoglobulin variable regions (Conticello 2008; Swanton et al. 2015). Therefore, the observed *APOBEC*-associated DNA editing signatures in birds are likely attributable solely to *A1*. Interestingly, passerines exhibit a markedly higher number of DNA editing sites and, with the exception of the Thraupidae order (*Camarhynchus parvulus* and *Geospiza fortis*), have retained *A1*. This pattern aligns with the reported adaptive expansion of ERVs in passerines (Chen et al. 2024). Although several avian *A1* genes show signatures of relaxed selection, the high ERV burden may explain why *A1* has been retained in these lineages.

## 5) Conclusion

There is a limited understanding of the evolutionary significance of *A1* across vertebrates, particularly in birds, despite its extensive study in mammals. Our findings reveal that *A1* has undergone a remarkably dynamic evolutionary history characterised by lineage-specific and independent inactivation events across all major bird orders, recurrent losses, and the emergence of an *A1*-like paralog unique to the Archelosauria lineage. The widespread pseudogenisation and loss of *A1* in birds, in contrast to its retention and diversification in other amniotes, suggest that selective pressures acting on *A1* have shifted substantially during avian evolution, likely in response to changes in ERV load and host defence mechanisms. Moreover, the lack of apoB RNA editing and the concurrent absence of apoE in birds suggest that the canonical lipid-regulatory function of *A1*, as seen in mammals, was lost early in avian evolution. Together, these results highlight *A1* as a model of functional plasticity—its ancestral role as a DNA editor repurposed for RNA editing in mammals but largely abandoned in birds. Further comparative and functional studies are needed to elucidate how the differential retention, duplication, and loss of *A1* and *A1*-like copies have shaped host genome defence, metabolic adaptation, and lineage-specific evolutionary trajectories across vertebrates.

## Ethics

This work did not require ethical approval from a human subject or animal welfare committee.

## Data accessibility

The data generated in this work, including sequence alignments, BLAST outputs, phylogeny and DNA edit sites, are available for download in the GitHub repository: https://github.com/AswinSSoman/APOBEC1_evolution_in_birds.git.

## Declaration of AI use

AI has been utilised to paraphrase sections of the text to enhance readability.

## Author contributions

A.S.S. wrote the manuscript with inputs from N.V. A.S.S. analysed all the data and wrote several scripts to automate important steps of the analysis. All authors reviewed the manuscript.

