## Supplementary Figures for "Deciphering *APOBEC1* in Avians: Unravelling loss events and functional insights"

### Contents

Supplementary Figure S1: Disruption of *A1* cis-acting factors in apoB RNA editing regions of birds……………….2

Supplementary Figure S2: Scatter plot of ERV content versus APOBEC1-associated G-A edit sites in birds………..3

Supplementary Figure S3: PacBio long read data mapped against the *A1* genomic region of *Struthio camelus*………4

Supplementary Figure S4: Nanopore long read data mapped against A1 genomic region of *Gallus gallus*…………..5

Supplementary Figure S5: PacBio long read data mapped against A1 genomic region of *Gallus gallus*……………..6

Supplementary Figure S6: Bionano optimal map data aligned to A1 genomic region of *Gallus gallus*……………….7

Supplementary Figure S7: PacBio long read data mapped against A1 genomic region of *Leptosomus discolor*……..8

Supplementary Figure S8a: Chromosome chain alignment of A1 and flanking genes of birds………………………..9

Supplementary Figure S8b: Chromosome chain alignment of A1 genic region of birds…………………………...…10

Supplementary Figure S9: Correlation between amount of DNA editing and *A1* loss…………………………….…..11

Supplementary Figure S10a: RNA-seq track of A1 and A1-like annotation region in as *Taeniopygia guttata*..…..…*12*

Supplementary Figure S10b: RNA-seq track of A1 and A1-like annotation region in as *Motacilla alba*……………..*12*

Supplementary Figure S11. Scatter plot of GC content versus G/C stretch length in the ApoE gene…………….….13

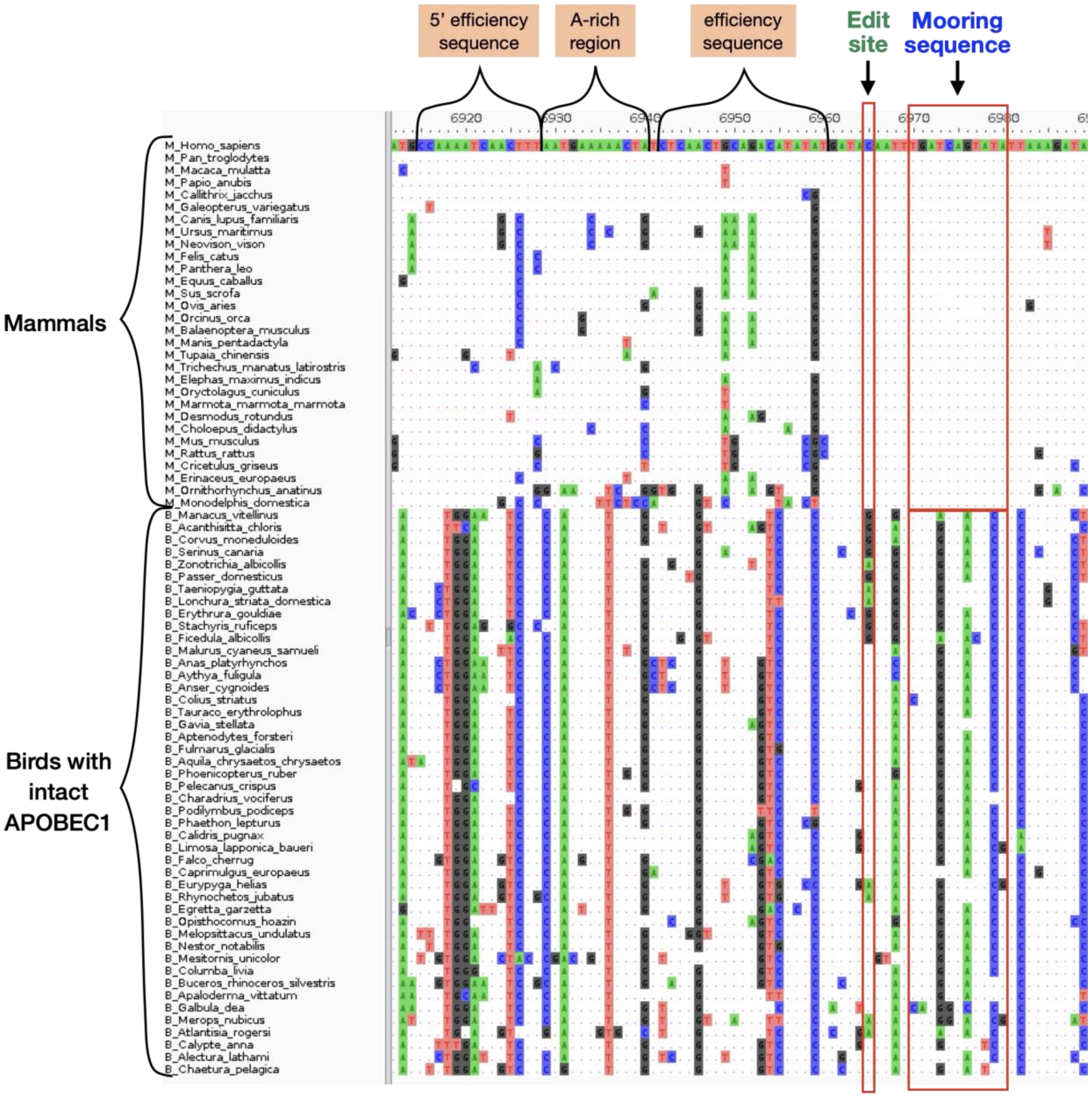


**Supplementary Figure S1: Disruption of *A1* cis-acting factors in apoB RNA editing regions of birds:** This figure presents a multiple sequence alignment of the apoB genic region, comparing cis-acting sequences required for apoB RNA editing in birds and mammals with intact *A1*. Dots indicate conserved sequences, and mismatches are highlighted in colour. Functionally relevant regions flanking the editing site are labelled, revealing numerous mutations throughout the alignment. The mooring sequence, a critical recognition element for target base editing, shows multiple mutations across bird sequences (red box), while this region remains highly conserved in mammals; some bird species also lack the edit site altogether. This pattern suggests that birds may be unable to undergo apoB RNA editing, despite the presence of a functional *A1* enzyme.

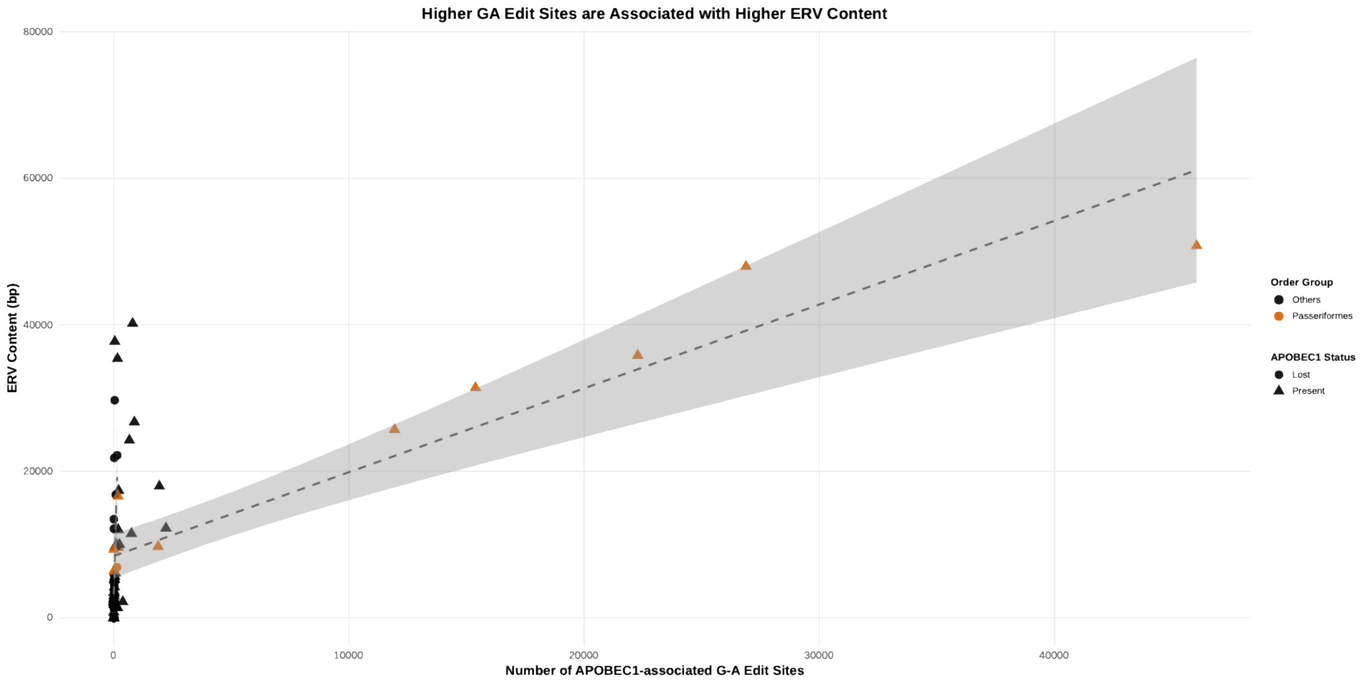


**Supplementary Figure S2: Scatter plot of ERV content versus *APOBEC1*-associated G-A edit sites in birds:** This plot shows the relationship between ERV content (X-axis) and the number of *APOBEC1*-associated G-A edit sites (Y-axis). Point shapes indicate the presence or absence of the *APOBEC1* gene, while colours distinguish bird orders such as Passeriformes (orange) and other clades (black). The dotted line denotes a positive correlation, indicating that a higher ERV load is associated with an increased number of *APOBEC1*-associated editing sites.

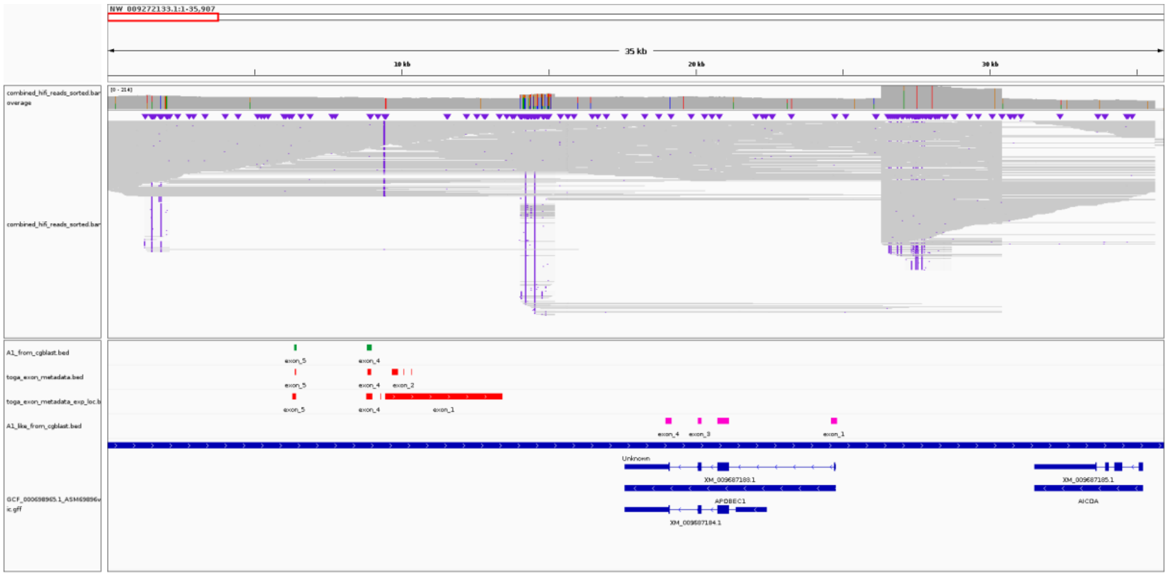


**Supplementary Figure S3: PacBio long read data mapped against the *A1* genomic region of *Struthio camelus*.** PacBio long reads were mapped from the assembly end to the downstream *A1* gene, *AICDA*. Although the assembly terminates before the upstream gene *NANOG*, the long reads span from the upstream region of the last exon (exon 5) of *A1*, as recovered from the assembly, extending to *AICDA*. The expected location of *A1* remnants is indicated by red bars, while the green bar shows recovered exons.

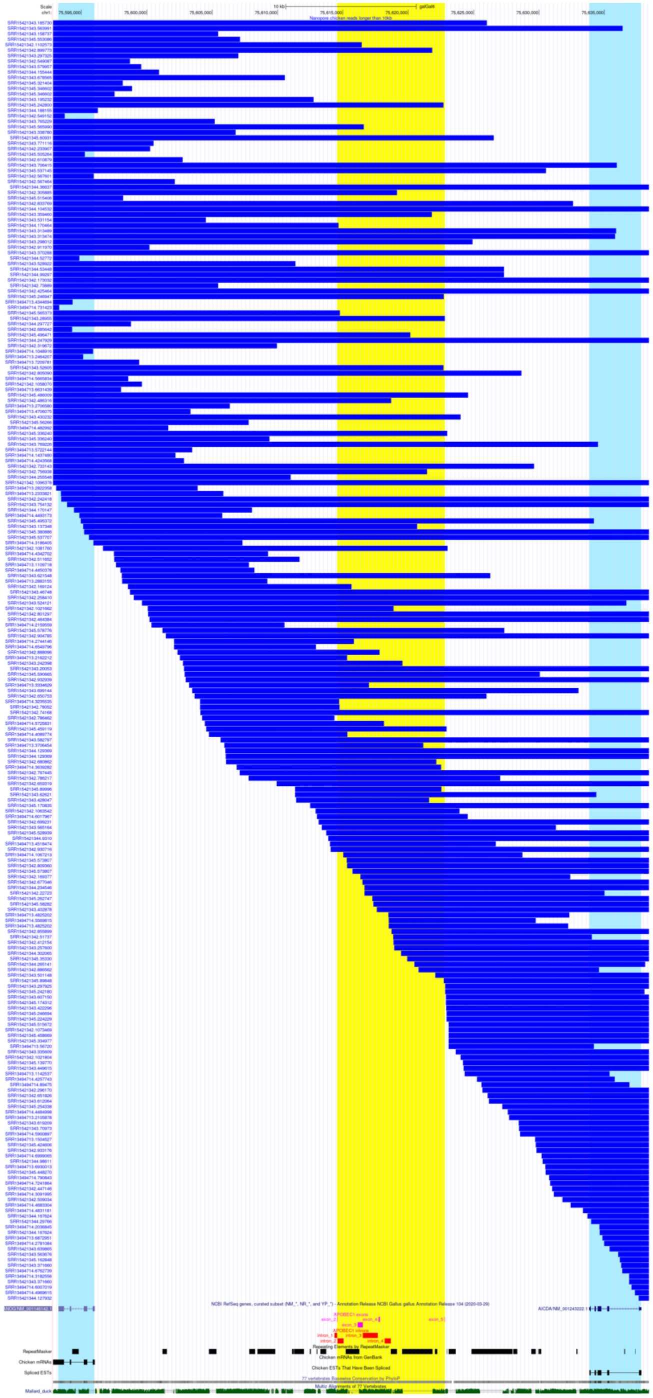

**Supplementary figure S4: Nanopore long read data mapped against the *A1* genomic region of *Gallus gallus*.** Reads longer than 10 kb were filtered, and the tiling path spanning the genomic region encompassing A1 remnants and the adjacent genes was visualised using the UCSC Genome Browser. Nanopore reads mapped to an approximately 45 kb region, spanning genes *A1*, along with upstream *NANOG* and downstream *AICDA*, show high coverage across these three genes.

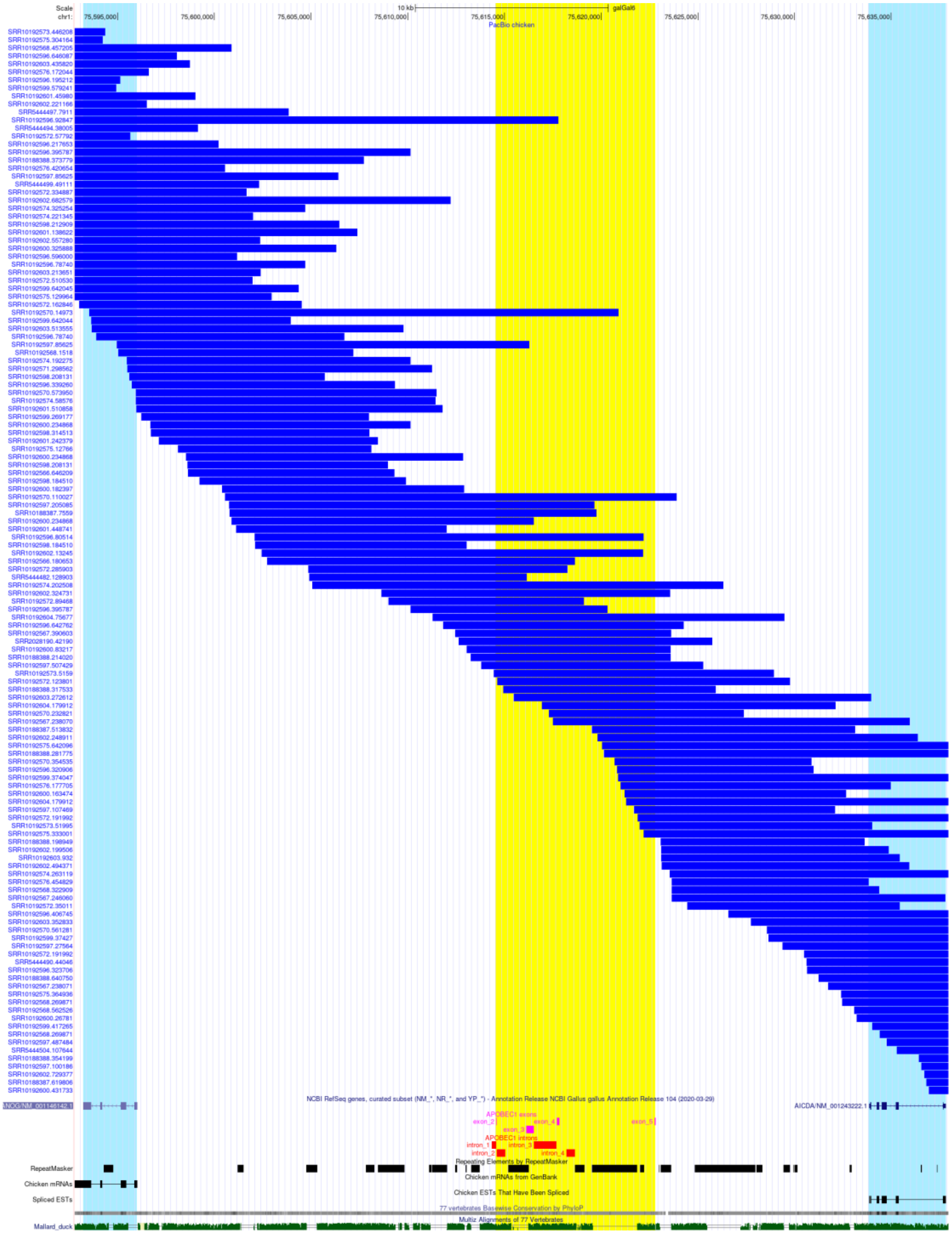

**Supplementary Figure S5: PacBio long read data mapped against the *A1* genomic region of *Gallus gallus*.** Reads spanning a region of more than 10kb encompass the genomic region containing *A1* and its flanking genes, without any drop in coverage, as visualised in the UCSC genome browser.

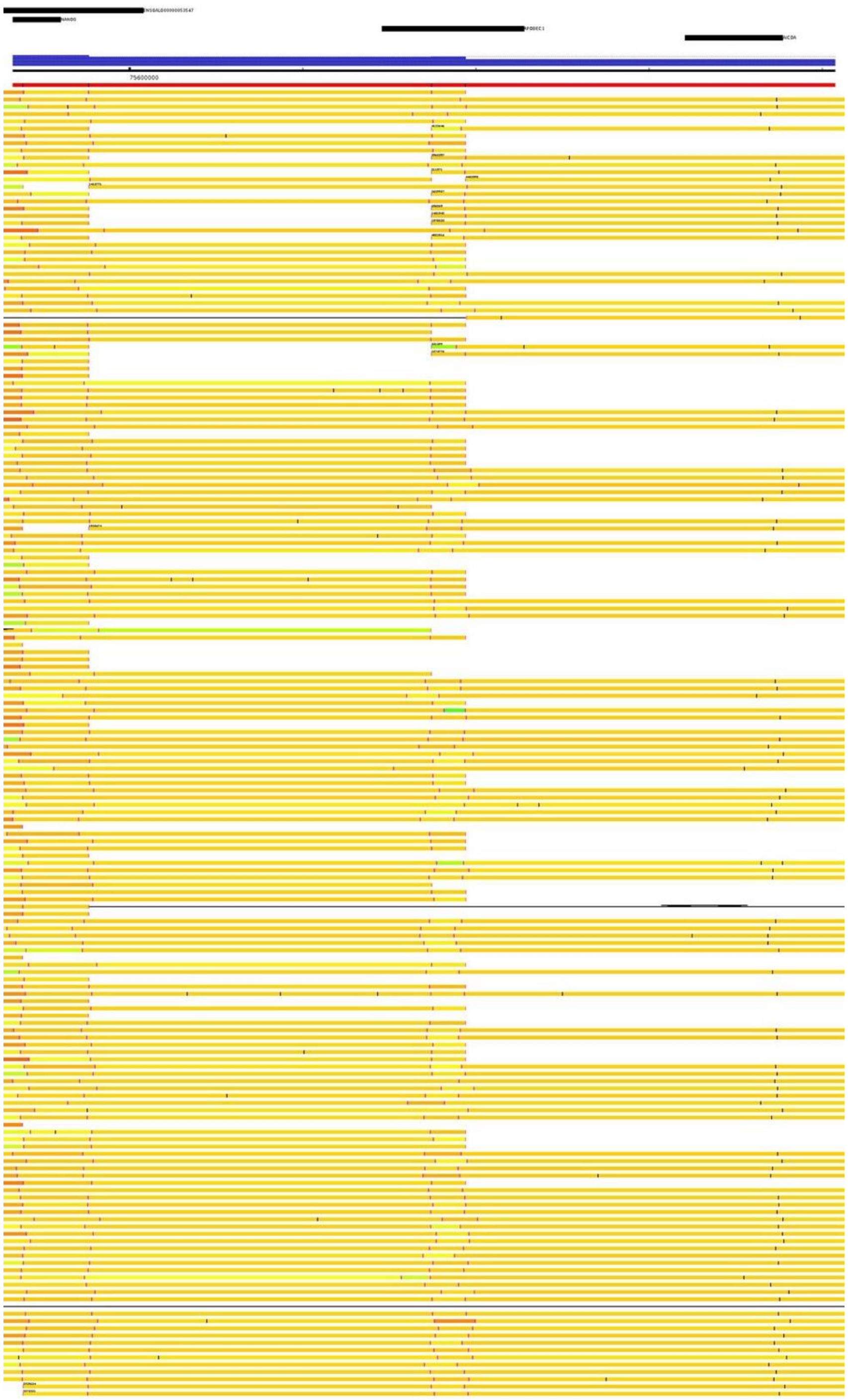


**Supplementary Figure S6: Bionano optimal map data aligned to the *A1* genomic region of *Gallus gallus*.** Optimal maps aligned to the region containing *NANOG*, *APOBEC1*, and *AICDA* display several single maps that span all three genes. All three optical maps (3273218, 3680518, and 3680519) spanned the *APOBEC1* region and validated the genome assembly.

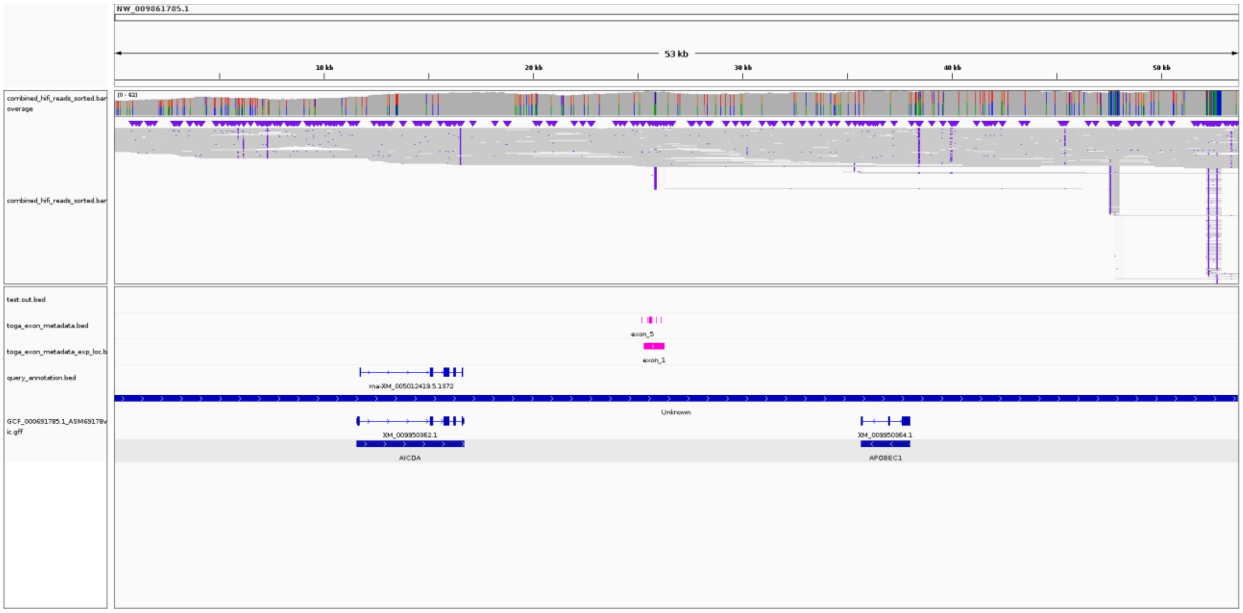


**Supplementary Figure S7: PacBio long read data mapped against the *A1* genomic region of *Leptosomus discolor*.** PacBio HiFi long reads show continuous coverage in the assembly, spanning the 8.2 kb deletion of *A1*, which removed all five exons up to the flanking gene *AICDA*. The expected location of *A1* remnants is indicated by magenta bars.

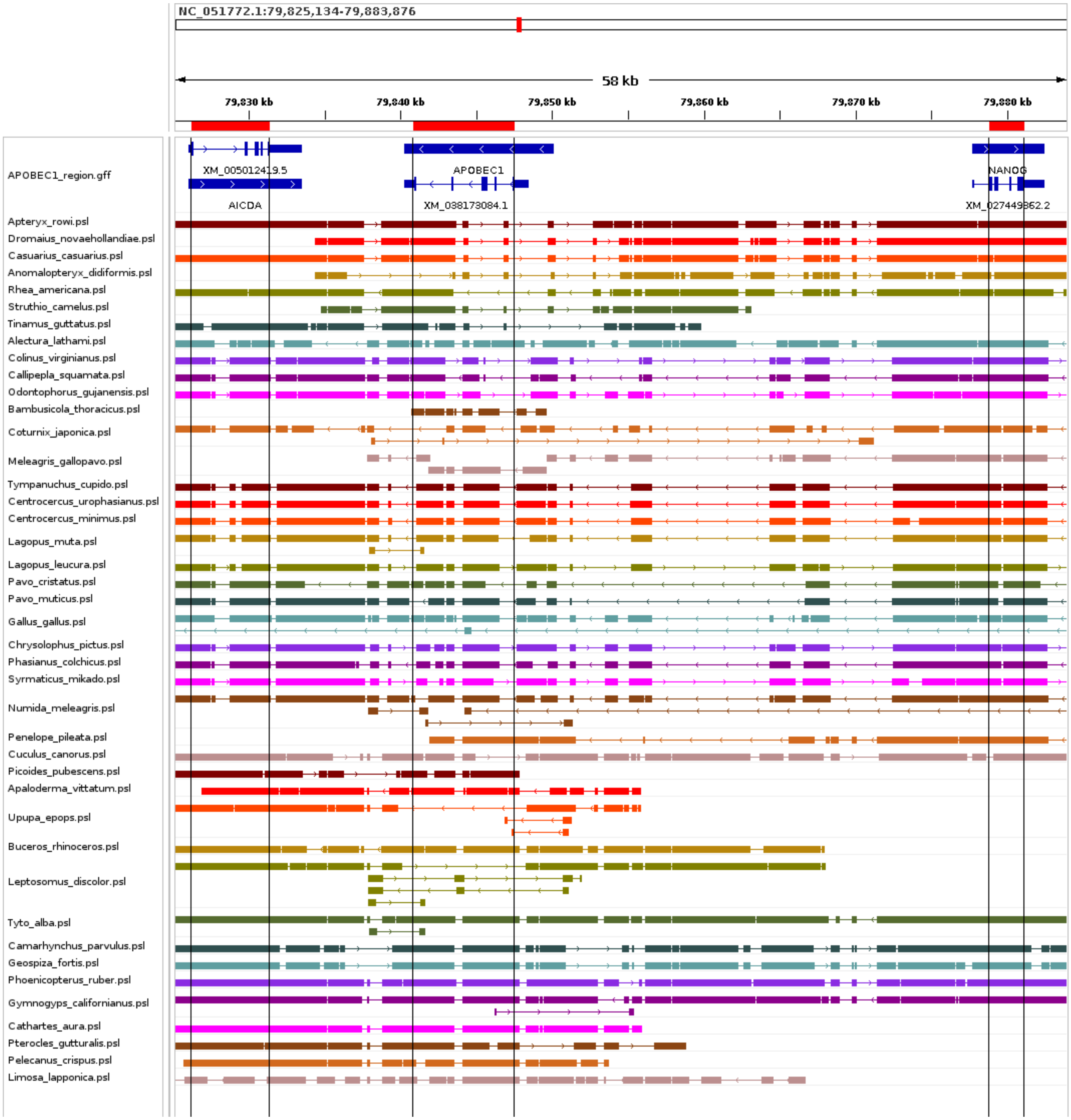


**Supplementary Figure S8a: Chromosome chain alignment of the region containing *A1* and flanking genes of birds.** Chromosomal chain alignments from the duck are shown as a reference at the top. Chains from each species are colour-coded. Black vertical lines with a red bar at the top indicate the coding region of each gene.

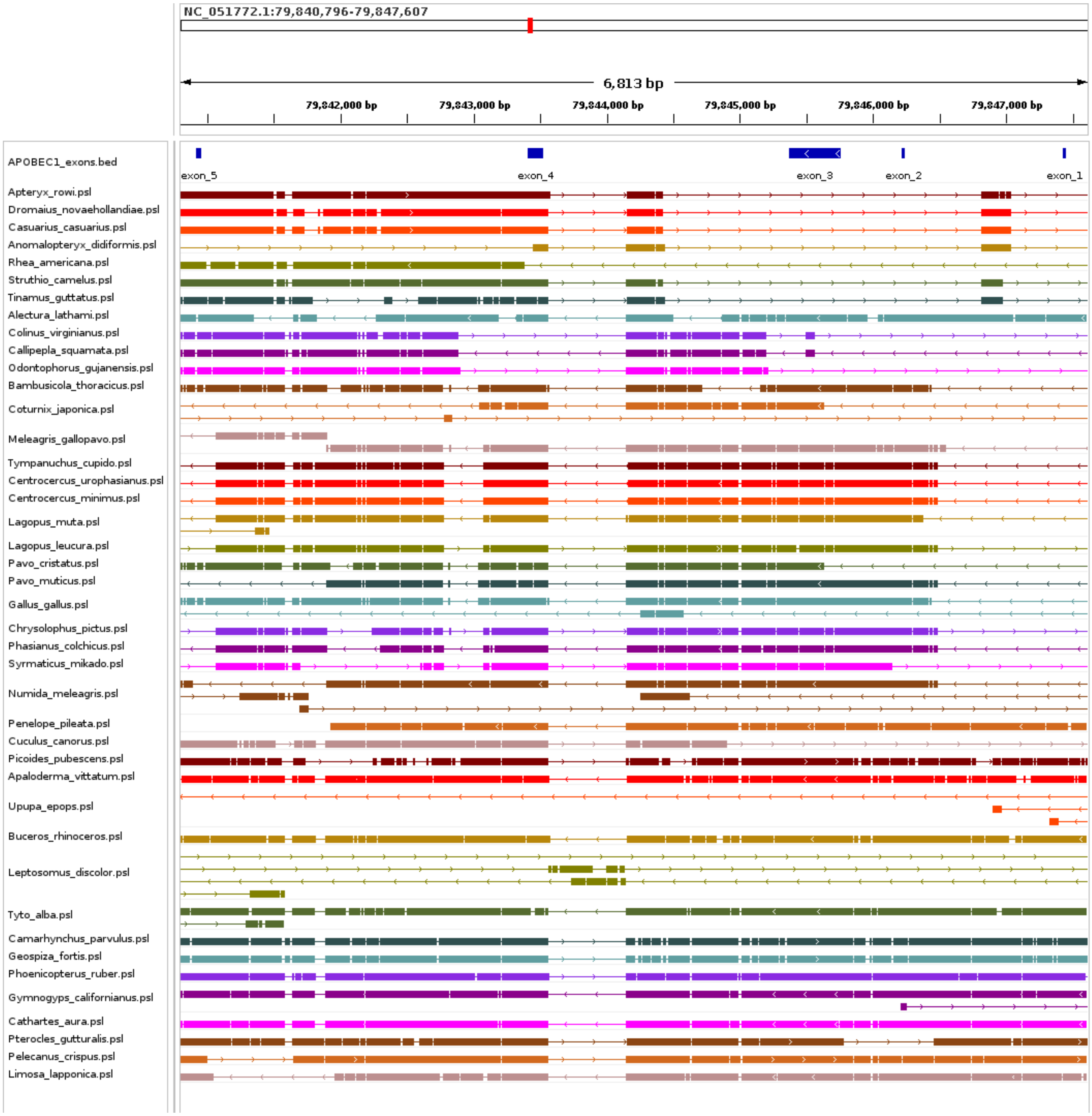


**Supplementary Figure S8b: Chromosome chain alignment of the *A1* genic region of birds.** Chromosomal chain alignments from the duck are shown as a reference at the top. Chains from each species are colour-coded.

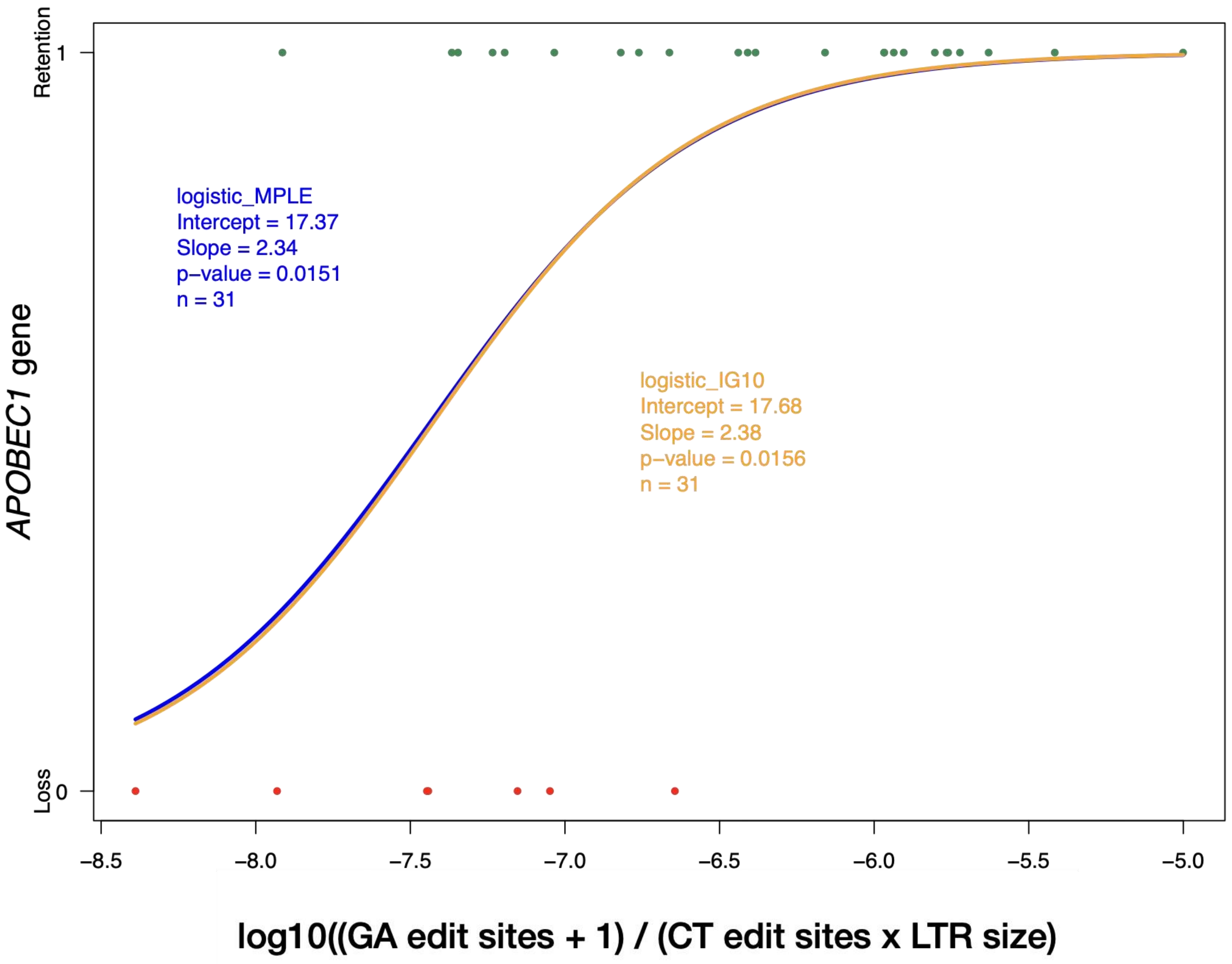


**Supplementary Figure S9: Correlation between amount of DNA editing and *A1* loss:** This figure illustrates the relationship between the presence or absence of *APOBEC1* (*A1*) and the abundance of *A1*-associated G-to-A DNA editing sites. To account for differences in genome size and background mutation rates, the number of G-to-A edits was log-normalised with respect to both the total LTR content in the genome and the number of C-to-T edits, which served as a negative control. The analysis reveals a positive correlation between *A1* presence and the abundance of G-to-A editing sites, with a statistically significant p-value. To evaluate the robustness of this association, two independent logistic fitting approaches—logistic MPLE and logistic IG10—were applied, both yielding consistent results.

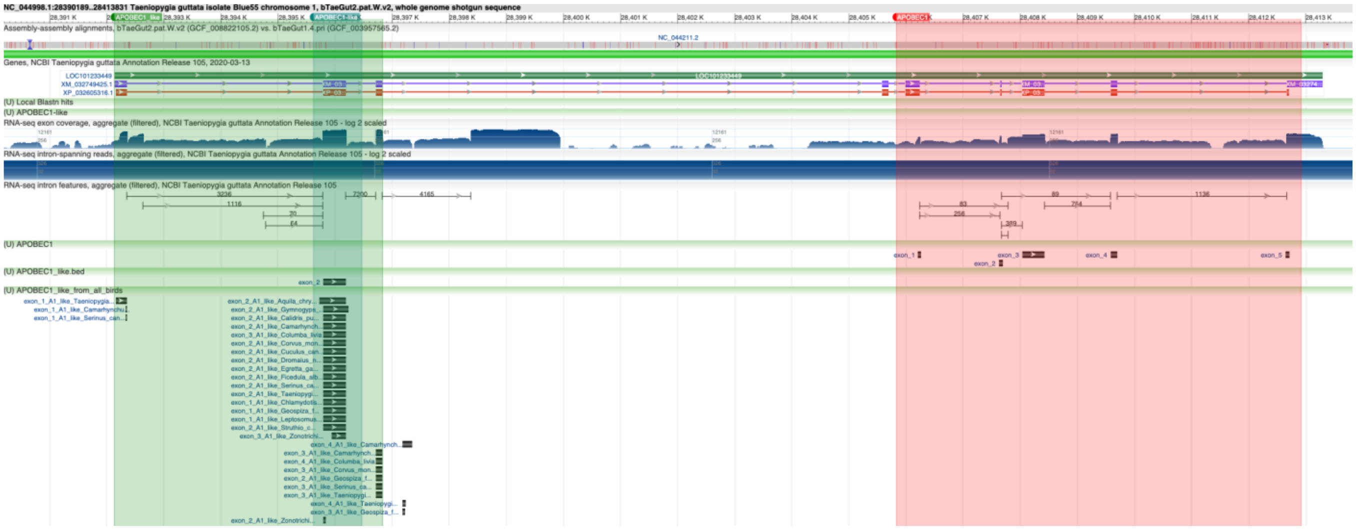

**Supplementary Figure S10a: RNA-seq track of *A1* and *A1*-like annotation region in *Taeniopygia guttata*:** RNA-seq track of *APOBEC1* of *Taeniopygia guttata* from NCBI genome viewer showing exon coverage and intron features.

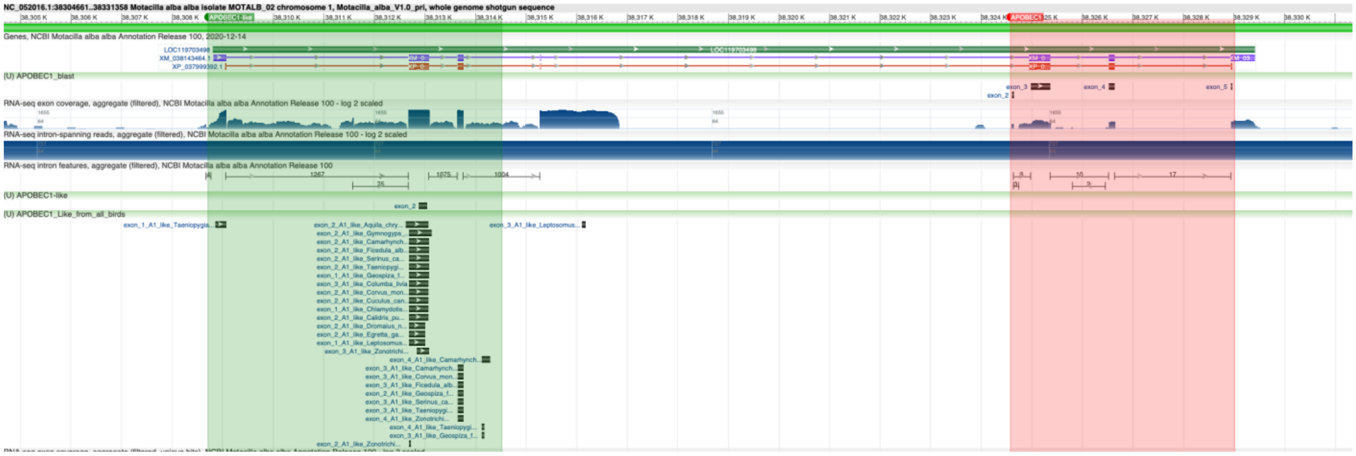

**Supplementary Figure S10b: RNA-seq track of *A1* and *A1*-like annotation region in as *Motacilla alba*:** RNA-seq track of *APOBEC1* of *Motacilla alba* from NCBI genome viewer showing exon coverage and intron features.

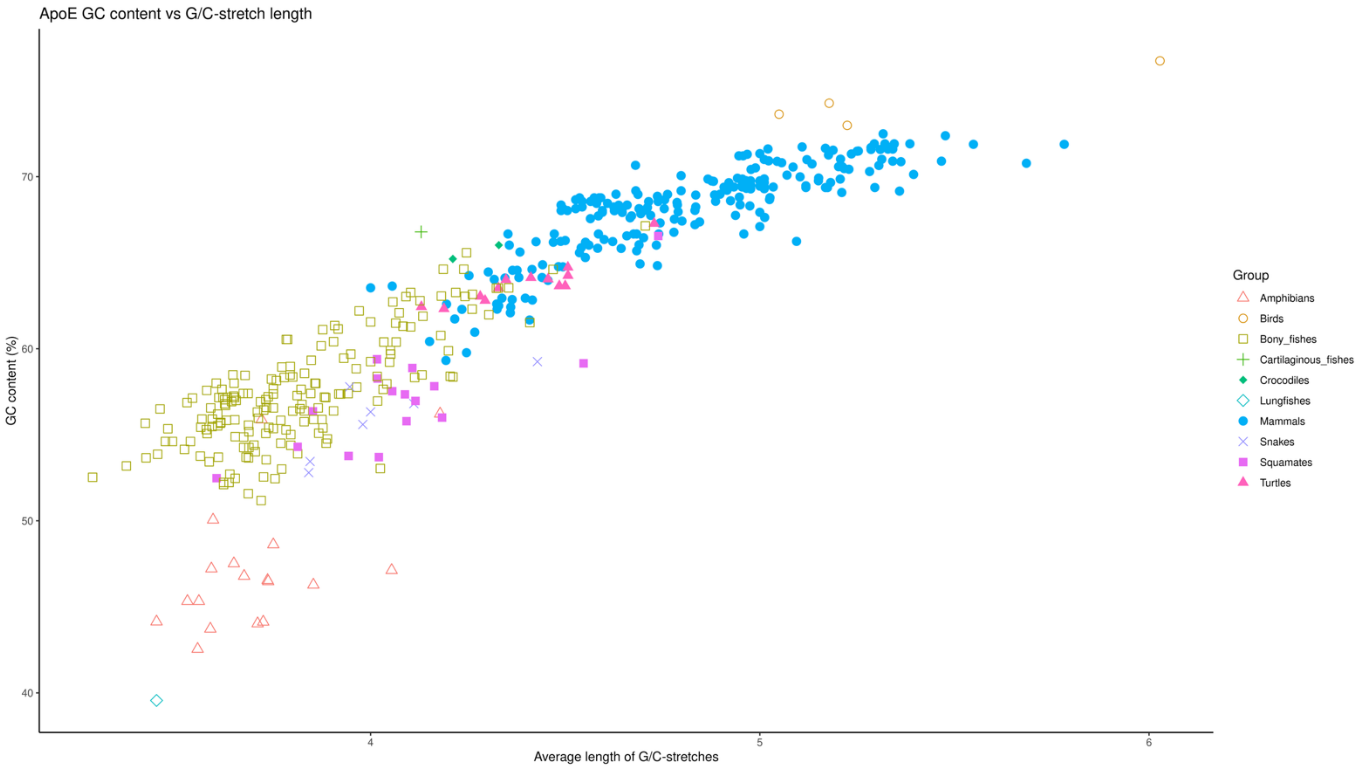


**Supplementary Figure S11. Scatter plot of GC content versus G/C stretch length in the ApoE gene:** This plot shows GC content on the X-axis and the average length of G/C stretches on the Y-axis for the ApoE gene. Point shapes and colours represent different vertebrate groups. The figure highlights that many mammals and birds exhibit high GC content in the ApoE gene, which can make accurate sequencing and recovery challenging.
